# StarPhase: Comprehensive Phase-Aware Pharmacogenomic Diplotyper for Long-Read Sequencing Data

**DOI:** 10.1101/2024.12.10.627527

**Authors:** James M. Holt, John Harting, Xiao Chen, Daniel Baker, Christopher T. Saunders, Zev Kronenberg, Nina Gonzaludo, Byunggil Yoo, Georgi Hudjashov, Maarja Jõeloo, James M.J. Lawlor, Weng Khong Lim, Estonian Biobank Research Team, Saumya S. Jamuar, Gregory M. Cooper, Lili Milani, Tomi Pastinen, Michael A. Eberle

## Abstract

Pharmacogenomics is central to precision medicine, informing medication safety and efficacy. Phar-macogenomic diplotyping of complex genes requires full-length DNA sequences and detection of structural rearrangements. We introduce StarPhase, a tool that leverages PacBio HiFi sequence data to diplotype 21 CPIC Level A pharmacogenes and provides detailed haplotypes and supporting visualizations for *HLA-A, HLA-B*, and *CYP2D6*. StarPhase diplotypes have high concordance with benchmarks where 99.5% are either exact matches or minor discrepancies. Manual inspection of the 0.5% mismatches indicates they were correctly called by StarPhase. With StarPhase, we update or correct 26.2% of GeT-RM pharmacogenomic diplotypes. Population distributions from StarPhase mostly reflect those of the All of Us cohort, while also highlighting gaps in existing pharmacogenomic databases that long-read sequencing can fill. With a single HiFi whole genome sequencing assay, StarPhase enables robust PGx diplotyping even as additional pharmacogenes and haplotypes are discovered.

## 1 Introduction

Pharmacogenomics (PGx) informs the safety and efficacy of treatments by providing predicted drug-gene interactions tailored to an individual genome. PGx applications range from broadly-applicable such as dosage and toxicity guidelines for anesthesia [11] to more focused guidelines such as viability of ivacaftor treatment for individuals with cystic fibrosis [6, 21]. Previous studies estimate 99% of individuals carry actionable PGx alleles [17, 4] and 20% of prescriptions have actionable drug-gene interactions [37], suggesting that PGx testing plays a critical role when prescribing drugs and therapies in personalized medicine. The first step of pharmacogenomics is identifying the haplotypes that are present within the patient, commonly termed PGx diplotyping. PGx diplotyping requires phased basepair-accurate genomic sequence, so haplotypes (referred to as star (*) alleles) can be identified using a predefined reference database.

The PGx diplotyping process can take many forms depending on the genomic complexity of the gene targets. The simplest pharmacogenes can be assessed by measuring the zygosity of a single genomic variant, such as rs9923231 in *VKORC1* [29]. However, most pharmacogenes require phased genetic variation across the gene, for example *CYP2B6* and *SLCO1B1* [29]. HLA genes are very diverse, so they are systematically defined with 4-field star alleles such that 2-fields define the protein sequence, 3-fields defines a cDNA sequence, and 4-fields defines a full-length DNA sequence [1]. *CYP2D6* is one of the most important pharmacogenes, but it is difficult to diplotype due to common structural variation including the *5 deletion allele, duplication haplotypes such as *2×2, and common hybrid alleles like *36 or *68 alleles [9, 34]. Duplications are often difficult to distinguish from single-copy haplotypes or masked by a deletion on the other haplotype, while hybrid alleles can be confused with the parent genes, *CYP2D6* and *CYP2D7*. Many existing assays provide partial solutions to these complexities, limiting the star alleles that can be assessed or restricting those alleles to abbreviated forms such as 2-fields for HLA genes.

The gold standard reference material data for clinical PGx testing, sourced from Genetic Testing Reference Materials (GeT-RM), was created from a consensus of nine labs using a composite of seven commercial assays, primarily microarray and PCR amplification technologies [28]. The GeT-RM PGx collection is a critical community resource for quality control, methods comparison, and validation. However, this collection has several known limitations including: 1) detection of only a subset of alleles, 2) inability to distinguish similar alleles, 3) inability to measure variant phase (i.e., in *cis* or *trans*), and 4) many caveats when assessing hybrid and copy-number changes in *CYP2D6* [28]. Additionally, new star alleles have been added to the PGx databases since this resource was created, many of which cannot be assessed from the previous assays and would require additional tests to update the collection. Short-read sequencing addresses some of these limitations, but the short reads rarely provide direct sequence evidence of variant phase, hybrid alleles, or copy-number changes. Many tools have been developed to perform PGx diplotyping using short-read sequencing [5, 20, 23, 31], reporting ambiguous or incorrect diplotypes when the short-read limitations manifest [13, 31, 32, 8]. Long-read sequencing can address many of the complexities with known pharmacogenes. In particular, HiFi sequencing provides accurate base-level sequence combined with long (15 kbp) read lengths [36] enabling direct observation of phased variants, full-length alleles, and copy-number or hybrid alleles. While some short-read PGx diplotypers have been adapted to accept long-read sequencing datasets, they often under-utilize the technology or produce errors [31, 14]. To our knowledge, there are no tools specifically designed to perform PGx diplotyping using only HiFi sequencing.

To address this gap, we introduce StarPhase, a new tool that utilizes HiFi sequencing to diplotype the 21 pharmacogenes from the Clinical Pharmacogenomic Implementation Consortium (CPIC [29]) where prescribing action is recommended (CPIC Level A): 18 variant-based haplotype definitions in the CPIC database and three complex genes (*HLA-A, HLA-B*, and *CYP2D6*) with definitions sourced from IMGT/HLA [1] or PharmVar [9] databases. We show that StarPhase has high concordance with comparator benchmarks (99.5% are matches or minor discrepancies), provides updated or corrected calls for 26.2% of GeT-RM diplotypes, and generates population distributions reflecting those of previous studies. StarPhase additionally provides full-length consensus sequences for *HLA-A, HLA-B*, and *CYP2D6*, enabling the reporting of detailed 4-field diplotypes for HLA genes as well as *CYP2D6* sub-alleles. These full-length sequences enable the discovery of alleles that are not described by the PGx databases, and the tool includes auxiliary metrics and visualizations that assist in the sequence-level verification of these potentially novel star alleles. StarPhase is easy to use, ingests standard inputs from typical HiFi processing workflows, and is available through both a pre-compiled binary and the source code.

## 2 Results

### 2.1 StarPhase accurately diplotypes pharmacogenes

StarPhase diplotypes 21 pharmacogenes using HiFi sequencing data and algorithms designed to leverage direct long-read sequencing observations. The input for StarPhase is a phased variant file (VCF), a phased alignment file (BAM), and a PGx database file (generated from [29, 1, 9]). The 18 genes sourced from CPIC [29] can be reliably diplotyped with only the phased variant calls from standard HiFi sequencing pipelines. For *HLA-A, HLA-B*, and *CYP2D6*, StarPhase generates consensus haplotypes from the alignment file and compares them to the database sequences. For *CYP2D6*, StarPhase utilizes the long reads to detect hybrid or duplicate alleles on the same haplotype. The primary output of StarPhase is a diplotype file, but it will also create full-length consensus sequences (FASTA) and detailed visualizations for *HLA-A, HLA-B*, and *CYP2D6*.

We first assess the accuracy of StarPhase on 147 datasets from the Human Pangenome Reference Consortium (HPRC, [35]) using a comparator set generated from multiple sources [31, 19, 5, 2]. These comparators were generated from either short-or long-read sequencing technologies and tools, and they collectively provide comprehensive coverage of all 21 genes reported by StarPhase. Of the 3,057 diplotype comparisons, 2,999 (98.1%) are exact matches with 15 of the 21 genes having 100% concordance across all datasets (see Figure 1). An additional 49 (1.5%) have minor discrepancies caused by differences in reporting or updates to the underlying databases. Included are 46 reporting discrepancies limited to two genes, *DPYD* and *RYR1*, that are caused by different protocols for handling haplotypes that are not defined in the CPIC database (see Supplemental Materials). The remaining three minor discrepancies are where StarPhase identifies haplotypes in *CYP2D6* that were added to the PharmVar database after the publication of the comparator set. Critically, these discrepancies are intentional, leading to a cumulative 99.7% of comparisons that are either matches or intentional discrepancies.

**Figure 1:**
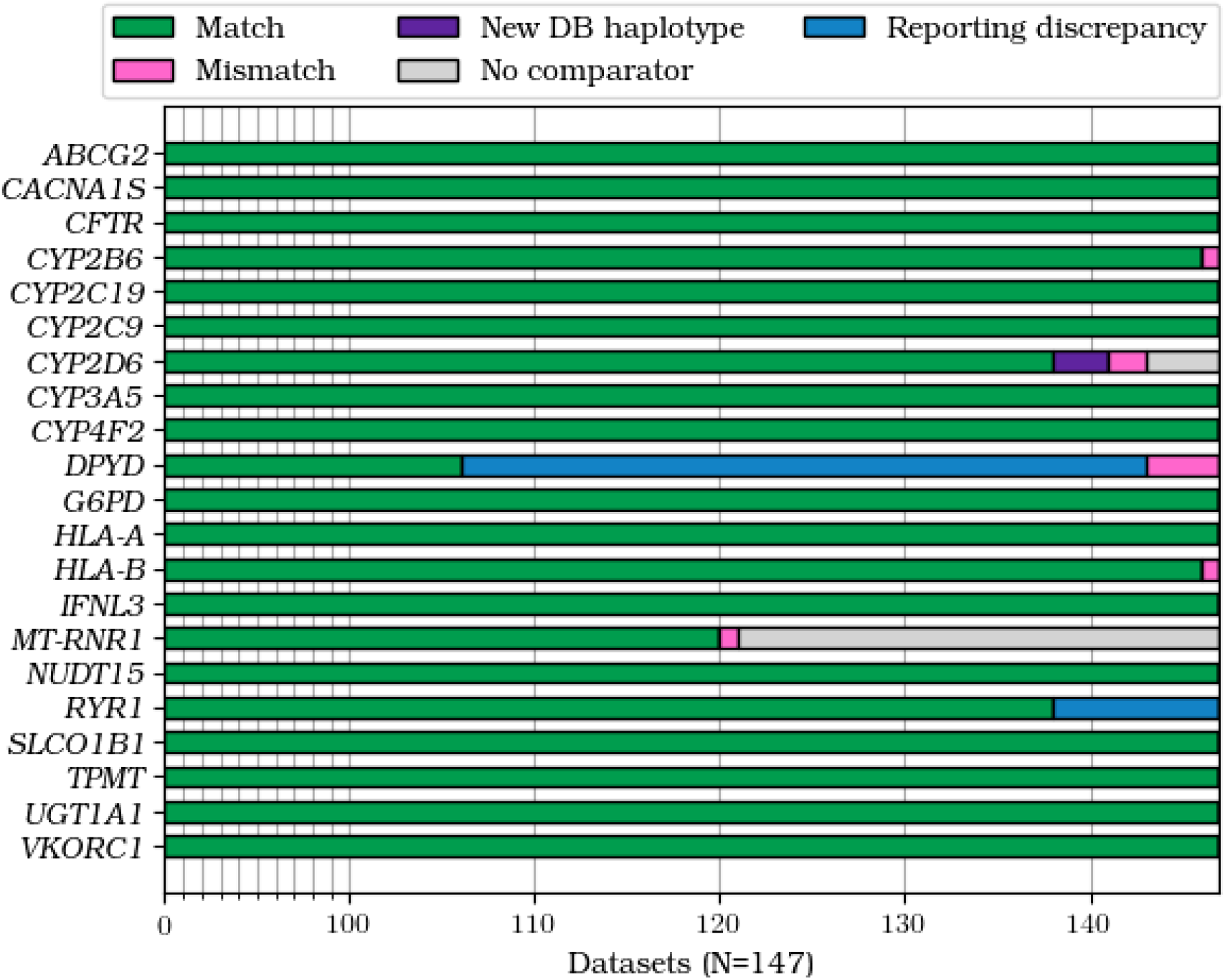
Summary of comparisons to HPRC comparator sets. For 147 HPRC datasets, we compare the diplotype calls of StarPhase to those from multiple comparator sets that collectively cover all 21 genes diplotyped by StarPhase. 30 diplotypes are missing a comparator value (“No comparator”), leaving a total of 3,057 comparisons. We identify 2,999 (98.1%) matches and 15 genes with 100% match rate. We observe three *CYP2D6* haplotypes that were called by StarPhase, but that were added to the database after the comparator set generation (“New DB haplotype”). There are 46 “Reporting discrepancies”, indicating comparisons where StarPhase and the comparator have different protocols for reporting diplotypes that are absent from the CPIC database. Lastly, there are nine total mismatches: one in *CYP2B6*, two in *CYP2D6*, four in *DPYD*, one in *HLA-B*, and one in *MT-RNR1*. Manual inspection supports that none of the mismatches or discrepancies are caused by diplotyping errors in StarPhase.

There are nine (0.3%) remaining “mismatch” comparisons with varying explanations from manual curation. The one mismatch in *CYP2B6* is caused by the comparator tool generating an error when it encountered an unexpected variant, leading to an unknown diplotype result. In contrast, StarPhase identifies the relevant variants and produces a diplotype call. The four mismatches in *DPYD* are likely errors in the comparator set caused by mis-interpretation of the phase information. The one mismatch in *MT-RNR1* is likely caused by different heteroplasmy allele fractions in the underlying datasets, leading to variable genotyping depending on the allele ratios in the sampled cell line. For the one mismatch in *HLA-B*, manual inspection supports the StarPhase diplotype call, but the source of the discrepancy is unknown. For *CYP2D6*, we believe the two mismatches are diplotyping errors in the comparator set: one likely caused by the presence of both duplication and deletion haplotypes in the same sample (see Figure 2) and the other with unknown origin. Critically, manual inspection did not identify any errors in the StarPhase diplotypes, and most of the mismatches appear to be errors in the comparator set (see Supplemental Materials).

**Figure 2:**
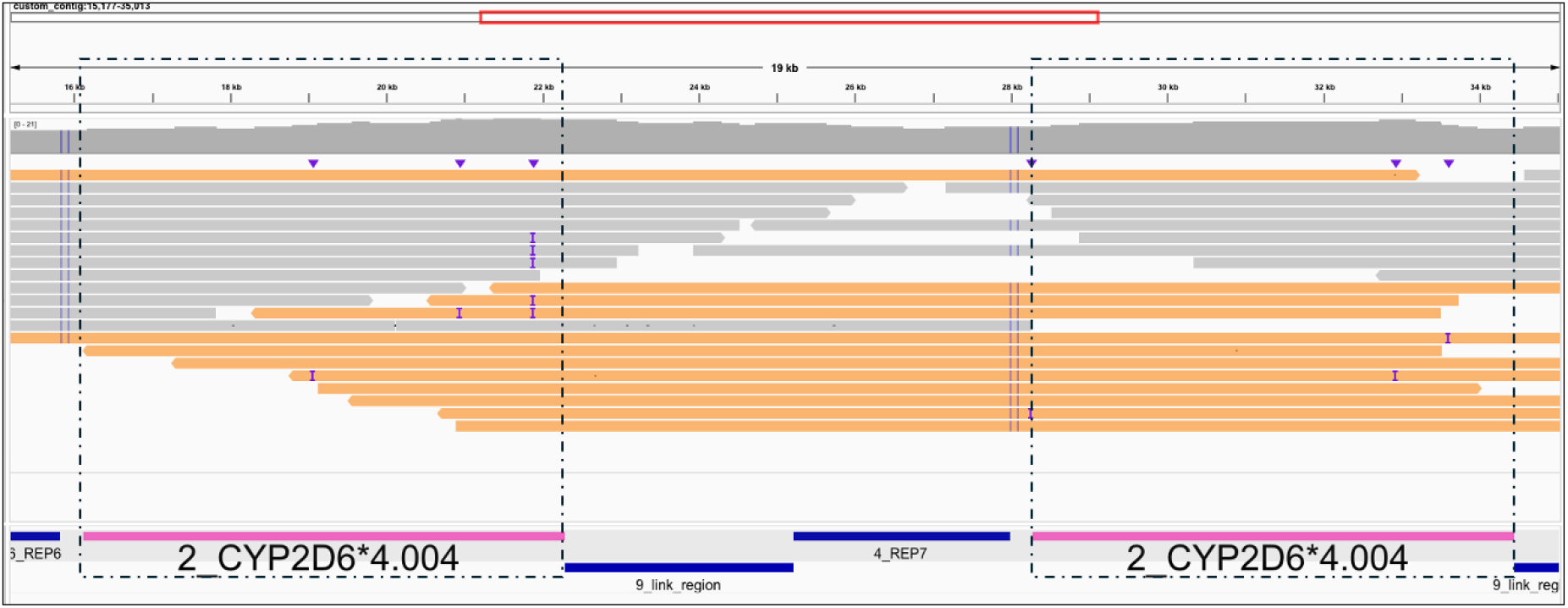
Example *4.004 duplication. StarPhase correctly identifies a *CYP2D6* *4.004×2/*5 (duplication and deletion) diplotype for NA20129. The comparator set for this dataset reports an incorrect diplotype of *4/*4, indicating that the deletion was obfuscated due to limitations of the underlying assay (short-read sequencing) and the duplication event. StarPhase generates multiple consensus sequences, assembles them into two sample-specific haplotypes, and then aligns the HiFi reads to these custom haplotypes to enable easier inspection through visualization software like IGV [30]. We show the portion of this assembled *4.004×2 haplotype with the duplication event, enhanced to emphasize the key evidence that is directly observable with HiFi sequencing. The two *4.004 alleles (pink BED regions with boxes) have identical sequence content with no observable consensus errors based on the re-aligned reads. We observe 12 reads (orange) that span both *4.004 alleles, providing direct sequence evidence of the duplication within this sample.

We perform a second assessment against the Genetic Testing Reference Material (GeT-RM) PGx benchmark [28]. As described earlier, this collection is the gold standard for validating clinical PGx pipelines, but it has many limitations including only detecting a subset of alleles, inability to measure variant phase, and difficulty distinguishing similar star alleles [28]. For our available datasets, this collection is restricted to only 12 of the pharmacogenes diplotyped by StarPhase. Of the 256 comparisons, there are 189 (73.8%) exact matches (see Figure 3). There are 38 (14.8%) StarPhase diplotypes with haplotype definitions that were not tested in the GeT-RM assays but visual inspection confirms. Additionally, the GeT-RM benchmark includes 22 (8.6%) diplotypes with at least one *UGT1A1* *60 haplotype, a haplotype which has since been retired from the CPIC database [24]. This indicates a combined 60 (23.4%) diplotypes where StarPhase is effectively providing updated, modern diplotypes for the GeT-RM samples (see Supplemental Materials).

**Figure 3:**
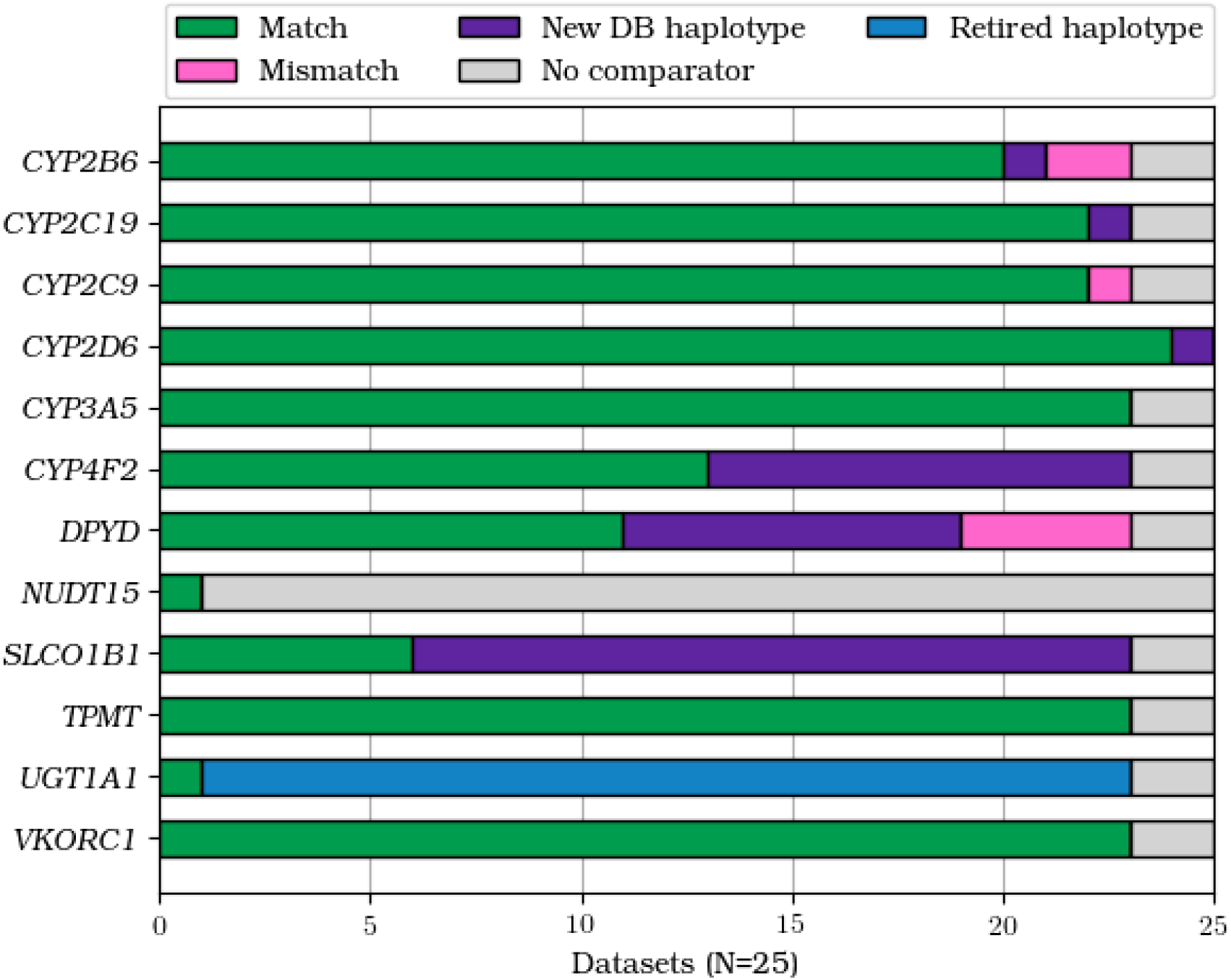
Summary of comparisons to the GeT-RM PGx benchmark. For 25 datasets included in the GeT-RM benchmark, we identify 256 possible diplotype comparisons. The StarPhase diplotype matches those from GeT-RM for 189 (73.8%). There are 38 (14.8%) comparisons where one or more haplotypes reported by StarPhase were not tested by the assays used to generate the GeT-RM diplotypes (“New DB haplotype”). Additionally, there are 22 (8.6%) GeT-RM diplotypes with *UGT1A1* *60, which has been retired from the CPIC database and is not reported by StarPhase (“Retired haplotype”). Lastly, there are 7 mismatches, 6 of which can be explained by database limitations or phasing errors in GeT-RM. We do not have a clear explanation for the last mismatch, but the StarPhase diplotype call is supported by manual inspection. In total, StarPhase provides updated or corrected diplotypes for 69 (26.2%) of the diplotypes in the GeT-RM PGx benchmark.

Lastly, there are seven additional mismatches. For five of these, StarPhase identifies haplotypes that do not match definitions in the CPIC database (*CYP2B6* and *DPYD*). For each of these, manual inspection reveals that StarPhase is correctly identifying the variants reported by GeT-RM, but with additional variants that were not tested by the GeT-RM assays and that do not form a defined database haplotype. We note that these are not errors from StarPhase, but are instead limitations of the existing databases definitions. The sixth mismatch is likely a phasing error in the GeT-RM comparator set caused by ambiguity in the assay (*CYP2B6*). For the last mismatch (*CYP2C9*), the StarPhase diplotype is supported by the underlying data, but the source of the discrepancy is unknown (see Supplemental Materials). Critically, manual inspection did not identify any errors in the StarPhase diplotype results, indicating that StarPhase is providing updated or corrected diplotypes for 69 (26.2%) of the GeT-RM benchmark diplotypes.

Across both comparator analyses, most of the comparisons are matches (96.2%). The most common sources of minor discrepancies are outdated comparators or reporting discrepancies caused by database limitations (3.3%), which have the greatest impact on assessments in *DPYD, SLCO1B1*, and *UGT1A1*. Importantly, manual inspection of all 16 (0.5%) identified mismatches support the StarPhase call, and the vast majority of these have an explanation for the discrepancy that was traced to the comparator sets. We provide all StarPhase diplotype calls from our accuracy assessment available as a comprehensive PGx reference resource for the community (see Supplemental Materials).

### 2.2 StarPhase diplotypes reflect known population distributions

We gathered 1,452 unrelated HiFi whole genome sequencing datasets from five contributing projects (see Supplemental Materials [7, 22, 15, 35, 3]). We ran StarPhase on each dataset and grouped the diplotypes into bulk metrics by ancestry and sex. We identify 441 distinct alleles across all genes in the cohort (see Figure 4). Three genes (*ABCG2, IFNL3*, and *VKORC1*) have only two defined alleles, each of which is observed in the cohort at least once. In contrast, *HLA-A* and *HLA-B* have the most diversity in our cohort, but also the smallest fraction of observed database alleles (*≤*2% each). Relatedly, some genes have low representation because the database haplotypes are exceedingly rare in the general population. For example, the two *CACNA1S* alleles that are not observed in our cohort are both non-reference alleles that were observed a combined 17 times out of 490,788 alleles (0.003%) in the All of Us cohort [12]. In total, the cohort includes 16,584 homozygous diplotypes, 11,036 heterozygous diplotypes, 675 hemizygous diplotypes (*G6PD* males), 1,433 homoplasmic diplotypes, 19 heteroplasmic diplotypes (1.3% of *MT-RNR1* diplotypes), and 475 diplotypes that did not match any pair of database haplotypes. The zygosity ratios vary by gene from 9.6% homozygous in *HLA-B* to 100.0% homozygous in *CACNA1S* (see Supplemental Materials).

**Figure 4:**
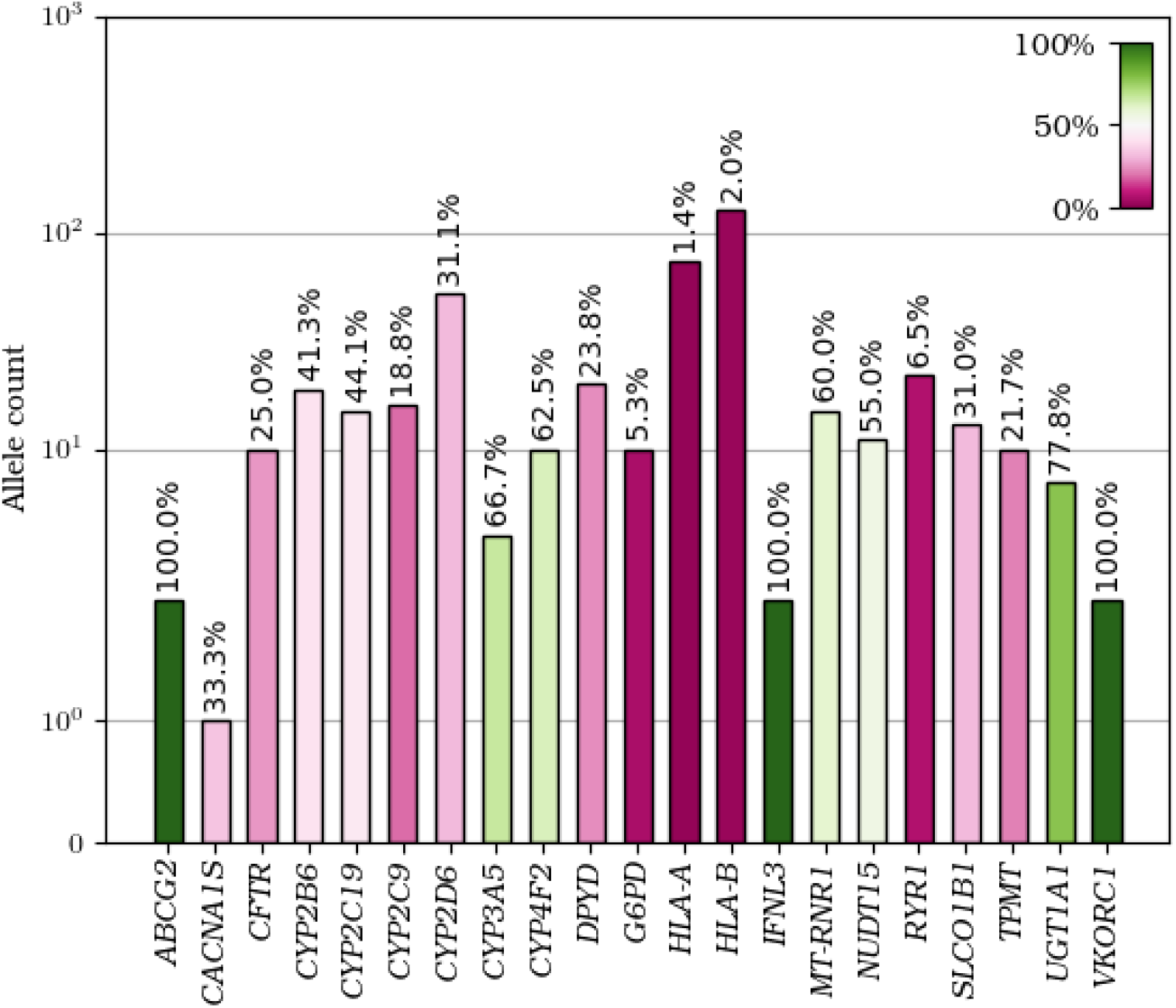
Diversity of alleles in StarPhase cohort. For each of the 21 genes reported by StarPhase, this figure shows the total number of distinct star alleles that are observed at least once in the cohort. Each bar is colored by percentage of the database, and these percentages are shown above each bar. *HLA-A* and *HLA-B* are reduced to their 2-field labels prior to counting (e.g., *52:01:02:01 and *52:01:02:04 both reduce to *52:01), and *CYP2D6* haplotypes are decomposed into individual core alleles prior to counting (e.g., “*4.002 + *68” counts as an observation of both *4 and of *68).

For the CPIC genes, we compare the relative haplotype distributions in our cohort to the population frequencies from the All of Us cohort [12]. Figure 5 shows the haplotype distributions of two genes, *VKORC1* and *SLCO1B1*, in both our StarPhase cohort and the All of Us cohort. In general, the relative percentages of the major haplotypes in our cohort are correlating with those from the All of Us cohort across all defined ancestries. For example, the “variant” haplotype of *VKORC1* is found at elevated levels in the East Asian (EAS) populations, decreased levels in the African (AFR) and South Asian (SAS) populations, and in-between levels in the American (AMR) and European (EUR) populations of both cohorts. The dominant haplotypes of *SLCO1B1* (*1, *37, *15, *14, and *20) are also correlated across all defined ancestries.

**Figure 5:**
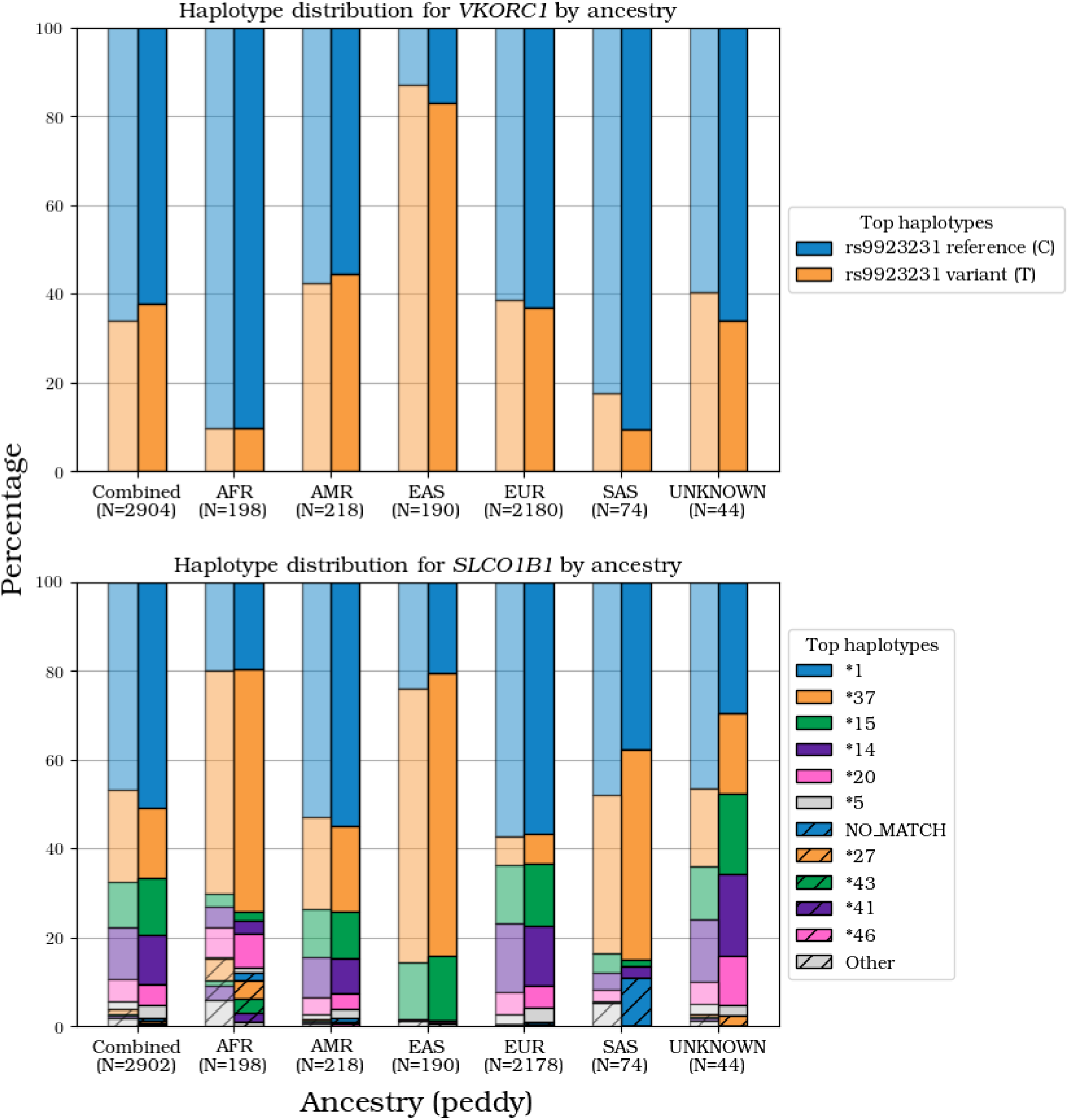
Haplotype distributions for *VKORC1* and *SLCO1B1*. These figures show the percentage of each observed haplotype in our cohort (solid right bars) and comparison percentages from the All of Us cohort (translucent left bars). Datasets are grouped by predicted ancestry, with “Combined” showing all datasets. The top haplotypes are ordered by their “Combined” frequency in our cohort. *VKORC1* is a relatively simple gene with only two haplotypes defined by a single variant. *SLCO1B1* is a more traditional pharmacogene with star alleles based on multiple variants. In both genes, the percentages of each major haplotype observed in our cohort are trending with the corresponding percentages from the All of Us cohort.

*CYP2D6* is the most complicated of the genes diplotyped by StarPhase due to the presence of deletions, duplications, and hybrids in the general population. Single-copy *CYP2D6* haplotypes are the most common type in our population, occupying the top 5 haplotypes in our cohort (see Figure 6). Haplotypes with an additional hybrid allele (*36 or *68) are the 6th and 8th most common haplotypes in our cohort, with the “*36 + *10” haplotype enriched in the East Asian population. The deletion haplotype (*5) is the 7th most common haplotype, and slightly enriched in the African population of our cohort. Duplication haplotypes are also observed, but these are relatively rare compared to the single-copy, hybrid, and deletion haplotypes. Figure 6 also shows the distributions of predicted diplotype metabolizer phenotype for our cohort. In general, the distributions match those previously described in [5], with “Normal” and “Intermediate” predicted metabolizer phenotypes as the two most often observed across all ancestries.

**Figure 6:**
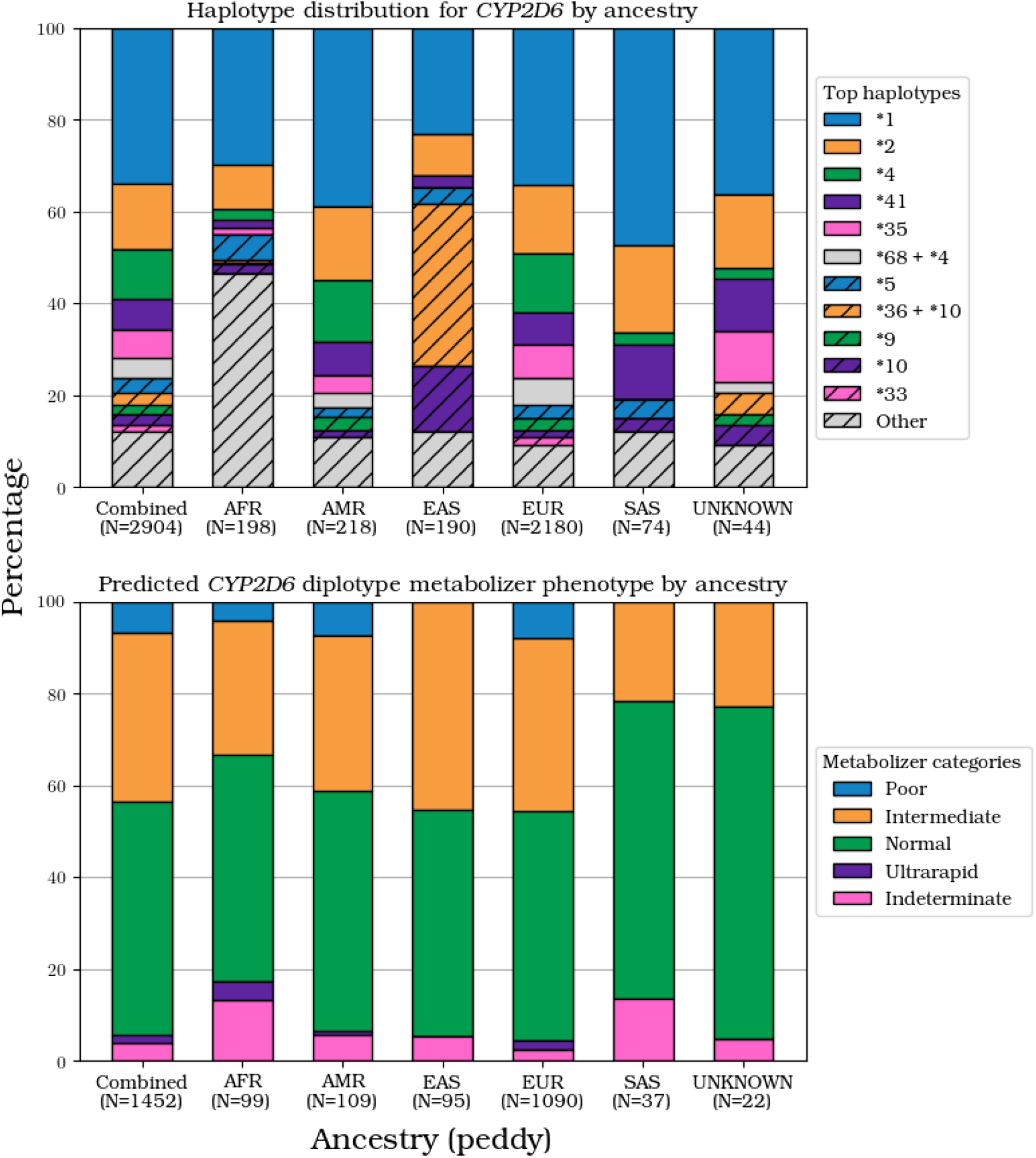
Population distributions for *CYP2D6*. These figures show the *CYP2D6* haplotype distribution (top) and the corresponding diplotype metabolizer phenotype distribution (bottom) by ancestry in our cohort. Haplotypes are ordered by their combined abundance in our entire cohort. StarPhase reports full *CYP2D6* sub-alleles from the database, but these are grouped by the core allele in this figure (e.g., *1.001 and *1.002 are reduced to *1). Most observed haplotypes have one copy of *CYP2D6* (top 5 haplotypes). Haplotypes with hybrid alleles (*36 and *68) are the 6th and 8th most common haplotypes in our cohort. Deletions (*5) are the 7th most common, while duplication haplotypes (e.g., *2×2 and *4×2) are relatively rare in our cohort (grouped with “Other”). Our cohort distribution of predicted metabolizer phenotypes matches those of previous studies [5].

Our cohort includes datasets where the observations do not cleanly match existing database definitions. For example, 475 diplotypes (1.6% of cohort diplotypes) do not have a pair of CPIC database haplotypes that exactly match the observations. This impacts 11 of the 18 CPIC genes, affecting over 1% of diplotypes for *DPYD* (22.8%), *RYR1* (4.7%), and *CYP2B6* (3.6%). *DPYD* is most affected because all current haplotype definitions are single-variant definitions, so any observed multi-variant haplotype has no matching definition in the database. Additionally, 1% of the *HLA-A* and *HLA-B* cDNA sequences did not exactly match the database, percentages that grow to 4% and 8%, respectively, when we consider the full DNA sequence. While some of these are probably technical artifacts, manual inspection reveals that many are likely haplotypes that are not described in the IMGT/HLA database (see Supplemental Materials).

## 3 Discussion

StarPhase is a comprehensive PGx diplotyping solution for HiFi sequence data. In our assessment against other sources, 99.5% of all comparisons are either matches or minor discrepancies, bolstering the accuracy of StarPhase. Manual inspection supports the StarPhase diplotype for the few mismatches (0.5%), and most of these have explanations for the difference revealing that StarPhase provides updated haplotypes, refines ambiguity, or corrects errors in the comparator diplotype calls. StarPhase reports detailed 4-field haplotypes for HLA genes, sub-alleles for *CYP2D6*, and full-length consensus sequences that enable basepair-level sequence resolution for some of the most difficult pharmacogenes.

We encountered several challenges with identifying accurate comparator diplotypes. Many existing assays are limited because they 1) test only a few star alleles, 2) cannot observe key distinguishing information such as phasing, full-length sequences, and/or copy-number changes, or 3) only test a subset of relevant pharmacogenes. This manifests as absent, ambiguous, imprecise, or incorrect diplotypes in each of our comparator sets. The greatest impact is in the GeT-RM PGx gold standard, which is primarily derived from microarray and PCR amplification assays, where StarPhase updated or corrected 26.2% of the diplotypes in our assessment. However, we also observe errors in our HPRC assessment which includes short-read sequencing based diplotypes. These limitations are largely resolved by HiFi sequencing, and StarPhase capitalizes on this advantage to provide accurate and detailed PGx diplotyping for the CPIC Level A pharmacogenes. We provide refined diplotype calls for all 21 genes reported by StarPhase for all datasets in our comparisons, creating a more comprehensive PGx reference for commonly-sequenced HPRC and gold standard GeT-RM samples.

While developing StarPhase, we identified some notable gaps in the existing PGx databases. In particular, there are several datasets where the observed haplotypes do not match any defined haplotypes in the CPIC database, with the greatest impact in *DPYD* and *RYR1* genes. These are typically caused by genes where all defined database haplotypes have a one-to-one relationship with a variant, leading to definition gaps for any observed multi-variant haplotypes. Recent joint consensus recommendations recognized this exact problem for *DPYD* and offered some practical clinical guidance for reporting [27]. However, these recommendations are unlikely to be broadly applicable to other pharmacogenes with similar multi-variant haplotype definition issues, and they do not address the core problem of undescribed haplotypes. We also observe multiple HLA sequences that do not exactly match those of the IMGT/HLA database, most of which are limited to DNA-only differences. Ideally, each of these undescribed haplotypes is systematically validated, annotated, and added to the corresponding PGx database through a rapid PGx annotation pipeline.

StarPhase is currently limited to the 21 CPIC Level A pharmacogenes. However, the methods can likely be extended to CPIC Level B genes and other pharmacogenes as they are discovered. StarPhase haplotypes are also limited to star alleles with definitions in the upstream PGx databases. Fortunately, the long reads are capturing surrounding variation that is not currently included in the PGx databases, and StarPhase is even identifying many haplotypes that are currently undescribed. In fact, the greatest source of discrepancy in our analyses came from the inability to describe those long-read observations using existing database haplotype definitions. We believe almost all of these discrepancies could be resolved with a database update that includes novel haplotype definitions matching our long-read observations. In other words, a single long-read whole-genome sequencing assay combined with StarPhase is robust to future database updates, enabling continuous re-analysis of PGx diplotypes without performing additional supplemental tests.

StarPhase is designed to leverage the accuracy and length of PacBio HiFi datasets for the purpose of pharmacogenomic diplotyping. Our initial gene set provides a comprehensive solution for CPIC Level A pharmacogenes. However, we believe that much of the underlying methodology in StarPhase is extensible to many other genes, if given appropriate haplotype definitions. For example, PharmVar has additional pharmacogenes that are not a part of CPIC Level A such as *CYP3A4*, and IMGT/HLA includes information on many more HLA genes including four additional HLA Class I genes. We plan to leverage the accuracy of StarPhase and HiFi long-read sequencing to provide additional diplotyping for these genes in the future.

## 4 Methods

### 4.1 Accuracy data gathering

The data used throughout our analyses reflects the realities of rapidly changing technologies and bioinformatic approaches in this field of research. While all datasets are from HiFi sequencing, these datasets were sequenced at different times with potentially different wet-lab procedures or sequencing platforms. The whole genome sequencing (WGS) and targeted sequencing datasets were pre-processed with publicly available workflows, but the exact versions of the underlying software may vary across datasets depending on when they were pre-processed. Each WGS dataset targeted *≥*30x coverage, whereas the targeted sequencing datasets tended to have well over 100x coverage within the target regions. The vast majority of both data collections were sequenced on the Sequel II platform. StarPhase was run on all accuracy datasets using a fixed version. Details on the bioinformatic workflows, tool versions, and tool commands are available in our Supplemental Materials.

### 4.2 HPRC comparator collection

We generated a comparator set collection for the CPIC genes using a version of PharmCAT [31] with a comparable database in terms of described haplotypes. We ran PharmCAT on the same collection of small variant VCF files that were provided to StarPhase, generating comparator diplotype calls for 17 of the 18 CPIC genes. Since we ran PharmCAT ourselves, all HPRC datasets that are a part of this analysis (N=147) were used for these 17 genes. The one CPIC gene that is not covered by PharmCAT is *MT-RNR1*, where we instead pulled comparator diplotypes from the 1000 Genomes Project (1KGP) [2].

For the HLA genes, we used a published comparator set that was generated using a combination of sequencing and bioinformatics technologies [19]. This comparator set includes 4-field diplotype calls for both *HLA-A* and *HLA-B* for 44 HPRC samples [19]. We identified 42 HPRC samples that were both in our data collection and this comparator set. For the remaining 105 datasets, we supplemented the calls by pulling results from the 1KGP HLA calls [2], each of which was limited to a 2-field diplotype. When comparing each HLA diplotype, we only required two fields to match (e.g., *52:01:02:04 would match *52:01:02:03) as those are most relevant to PGx downstream interpretation. We did identify five 4th field mismatches in *HLA-B* indicating a slight difference in detected DNA sequence, each of which is detailed in our Supplemental Materials.

Lastly, for *CYP2D6*, we used a comparator set that was published as part of the Cyrius paper [5]. This comparator set was generated by running Cyrius on short-read whole genome sequence datasets and includes notes on diplotype calls that failed trio validation or are likely incorrectly diplotyped. We identified 143 HPRC samples that were both in our data collection and this comparator set. These comparator *CYP2D6* calls did not include full sub-alleles, so we relaxed the match requirement to only require the core alleles match (e.g., *4.002 would match *4). Details on how we accessed or generated each comparator set is available in our Supplemental Materials.

### 4.3 GeT-RM comparator collection

We identified 25 total HiFi datasets (3 with WGS and 22 with targeted sequencing) that included at least one PGx diplotype call from the GeT-RM comparator set [28]. There were 12 total genes that were a part of this analysis, most of which had 23 datasets (1 WGS, 22 targeted) for our comparison. The two outliers are *CYP2D6* where we have 25 datasets and *NUDT15* where we only have one matching dataset. In contrast to the HPRC comparator sets, the GeT-RM collection is much older (published 2016, [28]) and includes haplotype labels that often do not match those of the modern CPIC or PharmVar databases. Where possible, we translated these labels to their modern forms prior to performing any comparisons. Additionally, the GeT-RM publication [28] includes a full list of tested haplotypes for each gene in addition to limitations of the tests, which we used to identify any haplotype calls generated by StarPhase that will never match the GeT-RM call because the haplotype was untested in the comparator assays. The GeT-RM comparator set also includes many imprecise diplotype calls. Our understanding is that this ambiguity comes from limitations of the technologies used to generate the call set. This occasionally led to automated parsing errors, followed by manual inspection of the two diplotype calls to determine if they were a likely match. Lastly, one haplotype, *UGT1A1* *60 was retired from the CPIC database [24], leading to discrepancies for almost all *UGT1A1* diplotype calls.

### 4.4 Population cohort data gathering

Our population cohort contains a mixture of datasets from rare disease projects [7, 15, 3] and population studies [22, 35], none of which were selected for pharmacogenomic reasons (i.e., we do not expect ascertainment bias). These datasets were inherently sequenced and processed under different conditions including: HiFi platforms (e.g., Sequel II, Revio), wet-lab procedures, sequencing depth, and bioinformatic processing pipelines. For secondary bioinformatic processing, each dataset required alignment to GRCh38 reference genome (e.g., pbmm2), small variant calling (e.g., DeepVariant [33]), and small variant phasing (e.g., HiPhase [16]) to generate the appropriate inputs for StarPhase. However, the exact tools, tool versions, and options vary across sites.

While the pre-processing of data was varied, we generated a standardized pipeline for our cohort collection that 1) ran StarPhase to generate diplotype calls for the 21 supported genes; 2) ran supporting tools to gather mean coverage [26], predicted ancestry, and predicted sex for each dataset [25]; 3) filtered out datasets with low mean coverage (*≥*15x required); 4) filtered out datasets that were computationally predicted to be 2nd-degree or closer relatives; and 5) collated the datasets into de-identified groupings based on predicted ancestry and sex. Importantly, only aggregate statistics from each contributing site were shared, preserving patient privacy for all underlying samples. We note that the relationship filtering step occurred at each contributing site. While it is unlikely that we collected relatives from different contributing sites, we cannot guarantee that there are no related individuals between the sub-cohorts.

Each contributing site downloaded this standardized pipeline and ran it locally on their datasets to generate de-identified sub-cohort aggregate statistics that could be shared. These sub-cohort aggregate statistics were then merged them into a single aggregate statistics file for our population analyses, grouping all results by computationally predicted ancestry and sex. The aggregate diplotype and haplotype call sets and a link to our standardized pipeline repository are included in our Supplemental Materials.

### 4.5 Population cohort analysis

The full cohort aggregate statistics file is the baseline data for population figures in this manuscript. *HLA-A* and *HLA-B* were reduced to 2-field diplotypes and *CYP2D6* reduced to core alleles for aggregate summaries. When comparing to the All of Us cohort, each defined ancestry correlated with one from the All of Us cohort. Our “UNKNOWN” ancestry was paired with the “oth” ancestry from the All of Us cohort.

For *CYP2D6*, predicted haplotype activity scores were derived using a lookup table available through PharmGKB [10]. The corresponding predicted diplotype metabolizer phenotype (Figure 6) was calculated by adding the two haplotype activity scores together and mapping the summation to the corresponding metabolizer ranges: Poor (=0.0), Intermediate ([0.25, 1.0]), Normal ([1.25, 2.25]), Ultrarapid (>2.5), and Indeterminate (anything with unknown or uncertain haplotype function) [18].

### 4.6 StarPhase core algorithms

StarPhase will generate diplotype calls for 21 different pharmacogenes. Of these, 18 are sourced from CPIC, two are HLA genes, and the last is the complex gene *CYP2D6*. The StarPhase database containing haplotype definitions is built from CPIC [29], IMGT/HLA [1], and PharmVar [9]. We briefly describe how each of these gene categories are diplotyped in StarPhase.

The 18 CPIC genes are all well-behaved from a long-read sequencing perspective, meaning that the read alignments and variant calls within the regions are usually high quality and trustworthy. StarPhase will extract all variant genotypes and phase information directly from the variant call file. One of the benefits of long reads is that the variants within a gene often reside within the same phase block, allowing StarPhase to accurately determine which PGx variants are in *cis* or *trans* for most of the CPIC genes. Importantly, StarPhase will correctly handle datasets with multiple phase blocks, which may generate ambiguity in the reported diplotypes. This occurs most frequently in longer genes such as *DPYD*. StarPhase will then compare the phased genotype calls to the reference CPIC database [29] and report all exact-matching haplotype pairs that support the observations. For the vast majority of cases, there is only one supported haplotype pair which StarPhase will report as the final diplotype for the gene. Additionally, if no supported haplotype pairs are identified, StarPhase will report the “NO MATCH” diplotype with a list of all identified variants in the detailed summary. This “NO MATCH” diplotype most frequently occurs in the CPIC genes that are primarily defined by individual variants as opposed to full-length haplotypes (e.g., *DPYD* or *RYR1*), and is the primary source of reporting discrepancy in our results.

In contrast to the CPIC genes, the diplotypes for *HLA-A* and *HLA-B* can not be consistently determined from a VCF file. False mappings of other homologous HLA genes to the wrong locus can generate false variant calls within the HLA regions and complicate diplotype calling directly from a VCF file. StarPhase resolves this by directly extracting mapped reads within each locus and discarding any reads that are erroneously mapped to the region. StarPhase then generates two consensus sequences (one for each haplotype) from the collection of remaining mapped reads using a dual consensus algorithm developed for this application (see Supplemental Materials). StarPhase limits the analysis to only reads that fully span each locus, allowing for highly accurate consensus sequences to be generated without ambiguity caused by partial overlaps. Each consensus sequence is then labeled using the IMGT/HLA database [1] (first by spliced cDNA sequence and second by full-length DNA sequence), and full-length 4-field diplotypes are reported when available. While the majority of our consensus sequences *exactly* match the sequences from the IMGT/HLA database, many of them do not. StarPhase can optionally report the generated consensus sequences with alignment metrics indicating the edit distance to the cDNA and DNA sequences for the reported alleles, enabling users to explore whether these deviations indicate an undescribed HLA haplotype.

Lastly, *CYP2D6* is the most complicated of the pharmacogenes diplotyped by StarPhase. The *CYP2D6* haplotype definitions include many complexities such as deletions (*5), duplications (e.g., *4×2), and hybrids with the nearby gene *CYP2D7* (e.g., *36 and *68). When any of the copy number or hybrid alleles are present, it becomes difficult to accurately diplotype *CYP2D6* using just a VCF file. StarPhase instead collects all reads that span the region and generates multiple consensus sequences from the read collection using a multi-consensus algorithm developed for this application (see Supplemental Materials). These consensus sequences represent either *CYP2D6* alleles (including hybrids), *CYP2D7* alleles, or the regions connecting them (e.g., REP6, REP7, or spacer region). StarPhase will then use the long-read observations to connect these regions together into putative “chains”, each of which may form a haplotype. All pair-wise combinations of chains are scored using a variety of heuristics, and the optimal chain pair is selected. Finally, the *CYP2D6* consensus sequences within the optimal chain pair are labeled using the haplotype definitions from PharmVar [9] and reported as a diplotype. For greater details on each of the diplotyping algorithms we implemented in StarPhase, we refer the reader to our Supplementary Materials and accompanying source code for StarPhase.

Supporting the above diplotyping algorithms, StarPhase also includes a variety of visualizations that readers may find useful for interpreting or verifying a StarPhase diplotype call. For the HLA genes and *CYP2D6*, StarPhase will map the inputs to the consensus algorithms back to the reference genome and then label them with their assigned consensus identifier. This is particularly useful for verifying accurate consensus clusters in the data through visualization software such as IGV [30]. These consensus sequences are also provided as debug outputs for review of possible novel, undescribed alleles. For *CYP2D6*, StarPhase will additionally generate the two full haplotype sequences as a custom reference genome and align all reads in the region to the custom reference. In addition to verifying consensus accuracy, this visualization helps provide direct visual evidence of duplication events that are often difficult to manually inspect in the GRCh38 reference alignments (see Figure 2).

## 5 Acknowledgments

Genomic Answers for Kids program is supported by generous gifts to Children’s Mercy Kansas City. Funding was provided by National Precision Medicine Programme (NPM) PHASE II FUNDING (MOH-000588) to W.K.L. Funding was provided by UM1HG007301, U01HG007301, R01HD112437, R01HG011598, Alabama Genomic Health Initiative (AGHI), and a grant from the Muscular Dystrophy Association (MDA 963255) to G.M.C. S.S.J. is supported by National Medical Research Council Clinician Scientist Award (NMRC/CSAINVJun21-0003). Funding was supported by SDDC/FY2021/EX/93-A147, 02/FY2022/EX/106-A168 and MOH-001005-00. Estonian Biobank calculations were carried out on the High-Performance Computing (HPC) Centre, University of Tartu, Estonia. The work done by M.J., G.H. and L.M. has received funding from the University of Tartu grant PLTGI23931 the European Union’s Horizon Europe research and innovation programme under grant agreement No 101060011. Views and opinions expressed are however those of the author(s) only and do not necessarily reflect those of the European Union or European Research Executive Agency. Neither the European Union nor the granting authority can be held responsible for them.

