## Supplemental Document for "StarPhase: Comprehensive Phase-Aware Pharmacogenomic Diplotyper for Long-Read Sequencing Data"

James M. Holt et al.

December 10, 2024

### 1 Data collections

#### 1.1 Tool and data versions

Table 1 contains all versions of each tool or dataset that was used in the analyses presented in this supplement as well as the main document.

#### 1.2 StarPhase database

The StarPhase database sources all information from other publicly available databases. This data is reformatted in preparation for diplotype calling in StarPhase. All database files are built using the `pbstarphase build` subcommand. These database files are periodically updated on the GitHub repository[1], but a user can generate a new one at their desired cadence. The following list contains details on how each database is accessed, versioned, and used in the StarPhase database:

1. CPIC database [19] - This database contains information on the 18 genes that can be diplotyped directly from a VCF file, which we commonly refer to as "CPIC genes" through this document. This database is accessed via the CPIC API (`https://cpicpgx.org/api-and-database/`). As of this writing, we are unaware of a way to access the CPIC version of the data via API, nor are we aware of a way to specify a particular version to download. Instead, the StarPhase database annotates the downloaded version based on a timestamp (e.g., `API-2024-07-30T14:45:20.368922802Z`). We note that while we refer to this as the "CPIC database" due to our API usage, many of the underlying haplotype and variant definitions are seemingly sourced from PharmGKB[2] and potentially other sources.

2. IMGT/HLA database [1] - This database contains full-length DNA and cDNA sequences for both *HLA-A* and *HLA-B*, as well as many other HLA genes that are not currently diplotyped by StarPhase. This database is accessed via the GitHub repository (`https://github.com/ANHIG/IMGTHLA`), which is also tagged with specific versions (e.g., `v3.57.0-alpha`). The version from the IMGT/HLA GitHub tag is propagated into the StarPhase database file.

3. PharmVar database [6]- This database contains the variants associated with *CYP2D6* haplotypes. This data is accessed by downloading the provided ZIP file on `https://www.pharmvar.org/gene/CYP2D6` and parsing out the various haplotypes. The version is implicit in the downloaded filename (e.g, `6.1.3`), which is then copied to the StarPhase database file.

4. Known constants - For *HLA-A*, *HLA-B*, and *CYP2D6*, StarPhase relies on knowing coordinates for various regions of interest (e.g., exons, introns, etc.). Additionally, there is prior knowledge for some genes that is also encoded in the database because it is not automatically captured by the other APIs (e.g., *CYP2D6* hybrid allele names). These values are encoded in the database file, and automatically populated with hard-coded constants when the `build` subcommand is called. We do not recommend naive users to alter these parameters, but they are available for experimentation and/or creating a database file for alternate reference coordinate systems (e.g., GRCh37, CHM13). The version for these coordinates is simply tracked via the full `pbstarphase_version` in the database (e.g., `0.13.0-3eff4a8`).

#### 1.3 Comparator call sets

The following list outlines where each comparator we used came from. The exact diplotype used for each comparison is copied into our Supplemental Tables.

1. CPIC gene comparators - These comparators were all generated by running PharmCAT [20] on the same VCF files provided to StarPhase, generating a diplotype for 17 of the 18 genes. The exact command is available in Section 1.5.

[1] Provided StarPhase database files: `https://github.com/PacificBiosciences/pb-StarPhase/tree/main/data`
[2] PharmGKB: `https://www.pharmgkb.org/downloads`

| Tool/Data | Version | Description |
|---|---|---|
| StarPhase | v0.14.2 | A phase-aware pharmacogenomic diplotyper for PacBio sequencing data that this document is about. Both the source code and a pre-compiled binaries are available on our GitHub repository. URL: `https://github.com/PacificBiosciences/pb-StarPhase` |
| StarPhase database | v0.14.1/20240826 | The database file provided with StarPhase containing CPIC, IMGT/HLA, and PharmVar haplotype definitions. Details on the contained data can be found in Section 1.2. This and other versions of the database file can be accessed through the GitHub for StarPhase. URL: `https://github.com/PacificBiosciences/pb-StarPhase/tree/main/data` |
| StarPhase cohort pipeline | v1.0.0 | This pipeline was shared with each contributing site to collect and aggregate PGx statistics based on de-identified ancestry/sex groups. It also contains scripts for generating the population figures from the main document and this supplement. URL: `https://github.com/holtjma/starphase_pipeline` |
| Data bundle | v1.0.0 | Contains the output files from our analyses. For the population cohort, this also includes a figure for each gene beyond what is highlighted in the main document. Zenodo URL: `https://doi.org/10.5281/zenodo.14289436` |
| WGS WDL pipeline | v1.x | This pipeline handles all data pre-processing for our WGS accuracy datasets. In short, it performs alignment (`pbmm2`), variant calling (DeepVariant [21]), and phasing (HiPhase [9]) on WGS HiFi sequencing datasets. The pipeline processed these datasets at different times with slightly different versions of the v1 pipeline. URL: `https://github.com/PacificBiosciences/HiFi-human-WGS-WDL` |
| HiFi target enrichment workflow | – | This pipeline handles all data pre-processing for our targeted sequencing accuracy datasets. In short, it performs alignment (`pbmm2` v1.10.0), variant calling (DeepVariant v1.5.0 [21]), and phasing (WhatsHap v1.4 [16]). Datasets were processed with different sequencing batches, and the exact version of the pipeline may vary between batches. URL: `https://github.com/PacificBiosciences/HiFiTargetEnrichment` |
| PharmCAT | v2.15.2 | Pharmacogenomic Clinical Annotation Tool, used to generate comparator calls for most CPIC genes. This version was the latest official version of PharmCAT when the StarPhase database used in our analyses was created. It was run via the Docker image: `docker://pgkb/pharmcat:2.15.2`. URL: `https://pharmcat.org` |
| *CYP2D6* function and phenotype references | Accessed 2024-08-29 | The "Allele Functionality Table" and "Phenotype Table" contain mappings from *CYP2D6* alleles to activity scores and then from total activity score (i.e., diplotype score) to predicted metabolizer phenotype. We accessed these reference tables through PharmGKB. URL: `https://www.pharmgkb.org/page/cyp2d6RefMaterials` |
| AllOfUs Biobank frequencies | v7, Accessed 2024-09-16 | AllOfUs cohort frequencies provided as part of [7]. We accessed the data through the "AllOfUs BioBank Frequencies" link on PharmGKB. URL: `https://www.pharmgkb.org/downloads` |

Table 1: Pipeline and data versions. This table contains a list of each tool or data file with the version used throughout this document. A description of each is provided, and direct URLs to each when available. For details on command usage for the tools, see Section 1.5.

2. *MT-RNR1* - For each sample, we pulled PGx results from DRAGEN v4.2.7 run on the 1000 Genomes Project [2] and parsed the corresponding *MT-RNR1* entry. The following is the URL pattern used for each {`sample`}: `https://1000genomes-dragen-v4-2-7.s3.amazonaws.com/data/individuals/hg38_alt_masked_graph_v3/{sample}/{sample}.final.star_allele.tsv`

3. *HLA-A* and *HLA-B* 4-field diplotypes - We downloaded a list of diplotypes provided by [11]. Specifically, we downloaded Supplementary Table 6 (`mmc6.xlsx`) and parsed out the relevant diplotypes for comparison, each of which was a full 4-field diplotype.

4. *HLA-A* and *HLA-B* 2-field diplotypes - For each sample, we pulled HLA results from DRAGEN v4.2.7 run on the 1000 Genomes Project [2] and parsed the corresponding HLA entries. These were all limited to 2-field HLA diplotypes. The following is the URL pattern used for each {`sample`}: `https://1000genomes-dragen-v4-2-7.s3.amazonaws.com/data/individuals/hg38_alt_masked_graph_v3/{sample}/{sample}.final.hla.tsv`

5. *CYP2D6* - We downloaded a list of diplotypes provided by [4]. Specifically, we downloaded Supplementary Tables S4 and S5 and parsed out the relevant diplotypes for comparison.

6. GeT-RM call set - We downloaded the file from GeT-RM[3]. Most of the calls we used are noted to originate from [18].

#### 1.4 Population cohort

Table 2 provides a summary of the sub-cohorts included in our population cohort. For each project, we provide a brief description of the purpose, sample types included, and the total number of samples in the sub-cohort.

#### 1.5 Tool commands

All tools were run via a Snakemake pipeline [10]. For the accuracy datasets, the command templates for each are included in the following sub-sections. For the population datasets, we refer the reader to the publicly available pipeline for each step (see Section 1.1).

##### 1.5.1 PharmCAT

PharmCAT is the comparator tool we used for most of the CPIC genes. This tool was run via Docker. Most of the following parameters specified input and output files. Of note, PharmCAT is only calling CPIC genes from a VCF file, so no additional reference or alignment files were required. The `--matcher-all-results` parameter will output all possible matches for a diplotype, matching the behavior of StarPhase when ambiguity is present. The `--missing-to-ref` parameter will interpret any entries that are missing as homozygous reference calls, matching a typical VCF file and the behavior of StarPhase. The command template for running PharmCAT follows:

```
/pharmcat/pharmcat_pipeline \
    -o {params.out_dir} \
    --matcher-all-results \
    --missing-to-ref \
    --reporter-save-json \
    --base-filename {wildcards.sample} \
    {input.vcf}
```

[3]GeT-RM PGx data page: `https://www.cdc.gov/labquality/get-rm/inherited-genetic-diseases-pharmacogenetics/pharmacogenetics.html`; File labeled as `Consolidated_GeT-RM_Table_20240418.xlsx`

| Project | Sample count | Description |
|---|---|---|
| Genomic Answers for Kids (CMKC) | 631 | A cohort of rare disease patient datasets included in the Genomic Answers for Kids (GA4K) program at Children's Mercy Kansas City [5]. The GA4K patients have a suspected genetic disease, and a large fraction of the patients are exposed to chronic drug therapy, including potential interactions with pharmacogenetic variation. The majority of GA4K patients represent primarily European ancestry as determined by Peddy analyses. |
| Estonian Biobank | 452 | The Estonia Biobank cohort contains a collection of datasets from adults in Estonia. Joining the cohort is voluntary, and there is no selection upon enrollment for disease or phenotype. The vast majority are of European ancestry as predicted by somalier [14]. |
| HudsonAlpha Pediatric Genomics | 154 | A cohort of probands and parents enrolled in one of several projects at the HudsonAlpha Institute for Biotechnology that perform genome sequencing on individuals with undiagnosed neurodevelopmental disorders [8]. ≈70% of probands are of European ancestry and ≈20% are African/African-American ancestry. Sequence data available via dbGAP/AnVIL, phs003537. |
| Human Pangenome Reference Consortium | 144 | A subset of the datasets used for accuracy assessment. Each sample is part of the Human Pangenome Reference Consortium [22]. These datasets are slightly enriched for African ancestry as predicted by somalier, but is otherwise fairly evenly split across ancestries. |
| Singapore Undiagnosed Disease Program | 71 | The Singapore Undiagnosed Disease Program is a multi-center, prospective cohort study of individuals with suspected genetic disorders [3]. The program was set up in 2014, and has recruited over 700 families (majority as trios) who have undergone a combination of targeted panel, whole exome, whole genome, transcriptome or long read whole genome sequencing. In conjunction with the genomic data, phenotypic data is available for all the patients. This includes demographics, medical diagnosis, laboratory, histopathological and imaging data. The majority of these datasets are predicted to have East Asian or South Asian ancestry. |

Table 2: Description of StarPhase cohort. This table outlines the projects that contributed to our cohort including a final filtered sample count and a brief description. All datasets are PacBio HiFi whole genome sequencing. Datasets within each project were filtered by coverage (requiring ≥15x). Additionally, datasets predicted to be 2nd-degree or closer relatives were filtered, leaving only unrelated datasets in each collection.

##### 1.5.2 StarPhase

StarPhase was run using the provided static binary file. The `--database` parameter is the database file provided through the StarPhase repository. The `--pharmcat-tsv` is a PharmCAT compatibile TSV file for downstream PGx associations. Currently, StarPhase only supports GRCh38 for the provided `--reference` file. All other files are standard input and output files.

```
pbstarphase \
    diplotype \
    -v \
    --database {input.database} \
    --reference {input.reference} \
    --vcf {input.vcf} \
    --bam {input.bam} \
```

| Gene | Matches | New DB haplotypes | Reporting discrepancies | Mismatches | No comparator |
|---|---|---|---|---|---|
| *ABCG2* | 147 | – | – | – | – |
| *CACNA1S* | 147 | – | – | – | – |
| *CFTR* | 147 | – | – | – | – |
| *CYP2B6* | 146 | – | – | 1 | – |
| *CYP2C19* | 147 | – | – | – | – |
| *CYP2C9* | 147 | – | – | – | – |
| *CYP2D6* | 138 | 3 | – | 2 | 4 |
| *CYP3A5* | 147 | – | – | – | – |
| *CYP4F2* | 147 | – | – | – | – |
| *DPYD* | 106 | – | 37 | 4 | – |
| *G6PD* | 147 | – | – | – | – |
| *HLA-A* | 147 | – | – | – | – |
| *HLA-B* | 146 | – | – | 1 | – |
| *IFNL3* | 147 | – | – | – | – |
| *MT-RNR1* | 120 | – | – | 1 | 26 |
| *NUDT15* | 147 | – | – | – | – |
| *RYR1* | 138 | – | 9 | – | – |
| *SLCO1B1* | 147 | – | – | – | – |
| *TPMT* | 147 | – | – | – | – |
| *UGT1A1* | 147 | – | – | – | – |
| *VKORC1* | 147 | – | – | – | – |
| **Total** | **2999** | **3** | **46** | **9** | **30** |

Table 3: Summarized comparison counts. For each gene reported by StarPhase, this table shows a summary of the comparison between StarPhase and the comparator diplotypes. This summary reflects the values visualized in Figure 1 of the main document.

```
    --output-calls {output.json} \
    --pharmcat-tsv {output.pharmcat_tsv} \
    --output-debug {output.debug_folder}
```

### 2 Discrepancies with HPRC comparators

Detailed notes on each discrepancy can be found in the accompanying Supplemental Tables (Section 5.1). Table 3 provides the exact counts of each match or discrepancy category for the figure from the main paper. The following sections provide broader descriptions of how we evaluated the identified discrepancies and categorized them.

#### 2.1 4th field discrepancies - *HLA-B*

For *HLA-A* and *HLA-B*, the bulk of the comparator haplotyped were limited to 2-field annotations, and 2-field is what is most critical for PGx diplotyping and downstream predictions. We chose to group all 2-field or better matches in the "Match" category for the main document. However, for 42 of our datasets, we had full 4-field diplotypes from [11].

When we use the full 4-field annotations, *HLA-A* was still an exact match for all 42 datasets, but we identified five discrepancies in *HLA-B*. Each of these differences impacted only one of the two haplotypes in each datasets (i.e., the other haplotype was a match) and each was constrained to a 4th field difference (i.e., the first three fields were a match). Sharing three fields of the HLA identifier indicated that they were identifying matching cDNA sequences, but that differences were detected in the full-length DNA sequence. Importantly, both 3rd and 4th field mismatches currently *do not* have any impact on downstream functional interpretation annotations as they are constrained to non-coding variation. 2-field matches are most important for downstream interpretation as changes there imply difference in the encoded protein which is more likely to impact function.

When we manually inspected each 4th field discrepancy, three of them shared a pattern where the consensus sequence generated by StarPhase was exactly 1 basepair edit distance from both the haplotype reported by StarPhase and the one in the comparator set (e.g., for HG01106 the consensus was 1 basepair edit distance from both *49:01:01:03 and *49:01:01:14). Additionally, when we inspected the corresponding aligned BAM file, the 1 basepair differences from each haplotype were supported by the reads. We believe these three are haplotypes that are not described in the IMGT/HLA database and that (by chance) are equi-distant from both the comparator haplotype and the StarPhase reported haplotype when compared using edit distance. For StarPhase, the metric for tie-breaking is the edit distance ratio (i.e., $edit\_distance/haplotype\_length$), so it will select the longer haplotype when the number of edits is identical. For the comparator set from [11], we are unsure if these ties were detected and, if so, how they selected one for the set. It is also possible that some of the newer haplotypes reported by StarPhase were not included in the comparator assays. Regardless, we would not categorize these discrepancies as errors in either StarPhase or the comparator set, but rather a limitation caused by the lack of exact-matching haplotypes in the IMGT/HLA database. We suspect that as the HLA database grows, these "ties" should become less frequent. However, users should be aware of these limitations, especially when assessing 4-field accuracy against any comparator set.

The fourth 4th field discrepancy is in dataset HG01928. When we inspected the consensus sequence, it was an exact match to the one StarPhase reported (*35:01:01:84) and had a 1 basepair edit distance to the comparator haplotype (*35:01:01:01). Manual inspection of the reads in the aligned BAM file also supported the StarPhase call. When we reviewed the database, we noticed this haplotype is relatively new to the database, added in August 2023 and updated in October 2023[4]. Given the overall high level of agreement between StarPhase and the comparator set, we suspect that this haplotype was simply not a part of the database used to build the comparator set (published in early 2024), and so the next closest haplotype was selected (*35:01:01:01). Regardless, the manual inspection supports that the *35:01:01:84 haplotype is the correct haplotype call for HG01928 given the current database entries.

The last 4th field discrepancy is likely caused by unequal representation of haplotype definitions in the IMGT/HLA database. For dataset HG01109, StarPhase reports *52:01:02:02 for one haplotype whereas the comparator set has *52:01:02:04. When we compare the consensus sequence to the DNA sequences for each haplotype definition, we identify edit distance values of 7 and 1 basepairs (bp), respectively. At face value, this would seemingly support the comparator haplotype (i.e., StarPhase making an incorrect call) and with a relatively large edit distance delta as well. However, these two haplotypes cannot be directly compared using just edit distance due to a significant difference in length of their database DNA sequences: 4,085 bp for *52:01:02:02 v. 2,725 bp for *52:01:02:04. One nuance of StarPhase is that it will first compare two potential HLA haplotypes within their *shared* alignment regions against the consensus sequence. In this case, the full length of *52:01:02:04 is contained within *52:01:02:02 and all the other bases are "extra" sequence that extends the haplotype definition upstream and downstream. When we manually inspected *just the shared region*, we found that the *52:01:02:02 haplotype *exactly* matched our consensus, whereas the 1 edit in *52:01:02:04 was located near the end of *HLA-B* and *inside* the shared region. In other words, all 7 edits for *52:01:02:02 are located in "extra" sequence relative to what was defined in the haplotype definition for *52:01:02:04. Additionally, manual inspection of the 1 edit relative to *52:01:02:04 did not appear to be caused by sequencing or processing errors, supporting StarPhase's *52:01:02:02 haplotype call within the shared region. In short, we believe StarPhase is making the correct haplotype call based on the shared region of these two haplotype definitions and the existing haplotype definitions from the IMGT/HLA database. We suspect that this haplotype is really a novel form of *52:01:02 that simply has not been described in the IMGT/HLA database yet. Figure 1 shows an IGV screenshot from the debug BAM file created by StarPhase for this dataset. We manually highlighted where each difference occurs between both the StarPhase haplotype and the one in the comparator set.

As mentioned earlier, this last discrepancy is mostly a problem with the IMGT/HLA database representations. These haplotype definitions have varying degrees of completeness in both the cDNA (e.g., some haplotypes have only a few exons instead of all eight) and in the DNA (e.g., the length and span relative

[4]Link to database entry: `https://www.ebi.ac.uk/ipd/imgt/hla/alleles/allele/?accession=HLA38088`

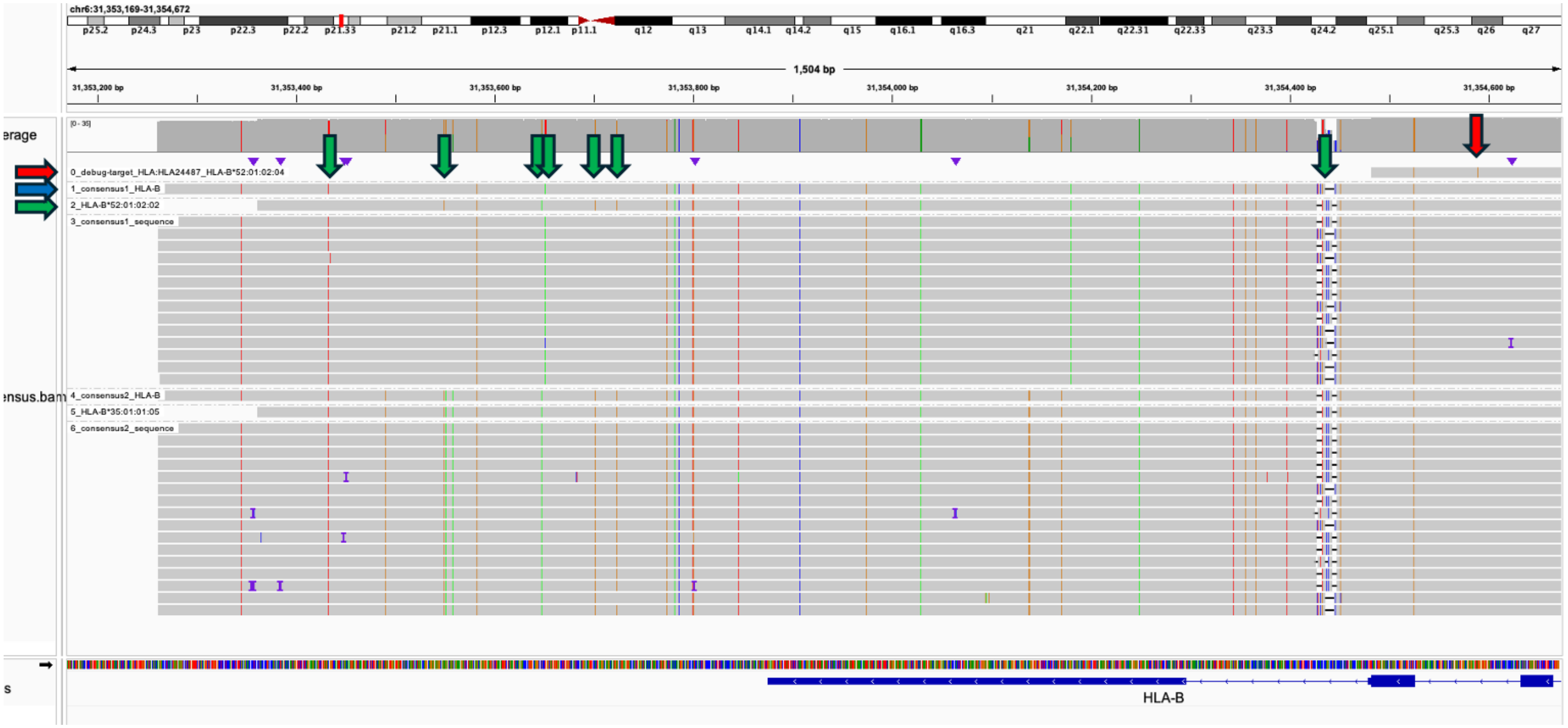

Figure 1: Example complex discrepancy in *HLA-B*. This figure shows an IGV screenshot from the debug BAM file created by StarPhase for one of our discrepant datasets. In this dataset, the comparator set indicated a *52:01:02:04 haplotype (sequence is along red horizontal arrow) whereas StarPhase indicated *52:01:02:02 (sequence is along green horizontal arrow). We found 7 edits between the consensus (sequence is along blue horizontal arrow) and the StarPhase call, but only 1 edit between the consensus and the comparator haplotype. However, when mapped to the reference genome, we can see that *52:01:02:04 is much shorter than *52:01:02:02 (2,725 bp v. 4,085 bp), creating approximately 1.3 kbp of "extra" sequence where additional edits can be found. In fact, all 7 edits between *52:01:02:02 and the consensus are found within this "extra" sequence (highlighted by green vertical arrows, 6 SNVs and 1 indel), whereas the 1 edit between *52:01:02:04 and the consensus is in the shared region (highlighted by a red vertical arrow). When the StarPhase algorithm compares the shared region from each of these haplotype alignments, the 1 edit for *52:01:02:04 is identified whereas *52:01:02:02 *exactly* matches the consensus within that shared region, ultimately leading to StarPhase reporting *52:01:02:02. While we believe the StarPhase call is more correct given existing database information, this problem is most likely caused by both a novel version of the *52:01:02 haplotype that is not described in the database and unequal representation of haplotypes within the IMGT/HLA database.

to the GRCh38 genes vary dramatically, as shown in our example). We suspect that enforcing greater stringency on the completeness of these haplotype definitions would alleviate this particular discrepancy, and potentially others that may be identified in the future. Additionally, we suspect that this haplotype is simply one that is not yet cleanly captured in the IMGT/HLA database (7 edits if we assume the StarPhase consensus sequence is correct or 1 edit if we assume the comparator is correct). We believe that improving the IMGT/HLA database by 1) enforcing some standardization on haplotype definitions and 2) further increasing the diversity of haplotype definitions in the database will help prevent future discrepancies.

#### 2.2 New database haplotypes - *CYP2D6*

When comparing the diplotype calls of StarPhase to comparator *CYP2D6* calls, we used published lists from [4], which was published in 2021. Since then, the upstream database has continued to grow as additional haplotypes have been identified in each of the genes. For *CYP2D6*, the exact list of alleles that can be identified by Cyrius can be found in the GitHub repository[5]. All 3 discrepancies that we have labeled as "New DB haplotypes" are absent from this list. Those alleles are *149.001, *152.001, and *155.001.

#### 2.3 Reporting differences - *DPYD* and *RYR1*

Most genes in the CPIC database have haplotypes that correspond to a collection of changes (i.e., multiple variants) with respect to the reference genome. However, there are many haplotypes that have one-to-one relationships between an identified variant and a haplotype definition. *DPYD* and *RYR1* are two such genes where this problem manifests frequently (discussed in the main paper population results). For example, the "c.3067C>A" haplotype is just a single variant change corresponding to c.3067C>A.

When StarPhase detects zero or one variants in a haplotype, the way to report the haplotype is understood. However, if two or more variants are detected in a haplotype, then the one-to-one relationships will prevent StarPhase from identifying any valid diplotype. Currently, StarPhase will report that there is no matching haplotype definition in the database ("NO_MATCH/NO_MATCH"). This is technically correct given the database haplotype definitions, but difficult to interpret downstream.

In contrast, the comparator tool (PharmCAT [20]) will allow for combinations of these variants into haplotypes that do not match a definition from the database. For example, if "c.3067C>A" and "c.46C>G" are identified on the same haplotype, it may report "c.46C>G + c.3067C>A" for that haplotype, despite not being part of the database definitions. These haplotypes are easily identifiable in the PharmCAT outputs by searching for haplotypes containing the "+" symbol in the output.

There are some issues with this approach, both from a definition and interpretation perspective. First, this method creates "novel" haplotypes that are not tied to a particular CPIC database haplotype. These haplotypes are unlikely to have any associated PGx interpretations, meaning they are not particularly useful for interpreting the impact on a patient. Second, while the community is starting to address this reporting problem for *DPYD* [17], there is still some ambiguity over how to report some of these haplotypes, especially when full-length phasing is unknown. Additionally, while many of the PharmCAT haplotypes generated this way are likely accurate, many of them are errors in VCF phase interpretation (see Section 2.4.3).

This problem was observed in our population cohort as well. In particular, *DPYD* (20.4%), *RYR1* (4.8%), *CYP2B6* (1.6%), and *SLCO1B1* (1.4%) each had over 1% of our cohort labeled with the "NO_MATCH" diplotype that could be traced to this issue. *DPYD* and *RYR1* are all defined by single variants in the CPIC database (one-to-one relationships), meaning there are *no* haplotypes where multiple alternate alleles are present. Despite having some defined multi-variant haplotypes in the database, this problem also impacts *CYP2B6* and *SLCO1B1*, where many one-to-one relationships between variants and haplotypes remain.

In the short term, we elected to keep the *DPYD* and *RYR1* process for haplotype identification identical to the other CPIC genes. For researchers wanting greater details, we note that the exact list of variants identified by StarPhase is part of the extended output JSON file. In the long term, we suspect that either 1) the CPIC database will be populated with multi-variant haplotypes as they are identified in datasets or 2) that the community will embrace a non-ambiguous, multi-variant haplotype reporting scheme which may be similar to structure PharmCAT has currently implemented. If the former is preferred by the community, then the existing StarPhase algorithms should automatically work as the database is updated. If the latter is

[5] Known Cyrius alleles: `https://github.com/Illumina/Cyrius/blob/master/data/star_table.txt`

preferred by the community, then we will likely alter the StarPhase algorithm in a future version to support this divergent reporting scheme.

#### 2.4 Mismatches

The following sections describe categorical reasons for mismatches on a per-gene basis. The exact set of identified mismatches can be found in the Supplemental Tables (see Section 5.1).

##### 2.4.1 *CYP2B6*

There was one diplotype called "Unknown/Unknown" by PharmCAT, where StarPhase called homozygous reference diplotype (*1/*1). In this dataset (HG03248), there is an unexpected heterozygous call: rs45482602, chr19:41009350C>[A, T]. While both alternate sequences have been found in population databases before, the CPIC database only includes the C>A version. This particular dataset was found to be heterozygous for the C>T version of the variant, leading to PharmCAT reporting the unknown result due to an unexpected variant sequence at the location. StarPhase was only searching for the C>A version (ignoring the other ALT sequence) and assumes homozygous reference if it does not detect that allele, ultimately leading to the homozygous reference diplotype call. In this instance, we do not categorize either tool as "incorrect", but the identification of unexpected alleles is handled differently.

##### 2.4.2 *CYP2D6*

There are two mismatches we identified in the *CYP2D6* that cannot be explained by new database haplotypes. The first dataset, "HG02492", is labeled as *106/*2 in the Cyrius set but was called *1.001/*2.001 by StarPhase. There is only one variant distinguishing *106 from *1.001 (rs28371733, chr22:42126914C>T). We manual inspected the aligned BAM file, but we saw no evidence of this variant in any of the reads. We then downloaded a short-read dataset for HG02492 from 1KGP[6] and re-ran Cyrius on this data. This re-run also produced a *1/*2 call, matching what was reported by StarPhase for the long-read dataset. We are unsure of why the Cyrius set has *106/*2, but we suspect it was either an error produced by Cyrius or perhaps a documentation error for that dataset. Regardless, we believe the *1.001/*2.001 call produced by StarPhase is correct.

The second data, "NA20129", is easier to explain. Here, the Cyrius diplotype is *4/*4 and the StarPhase diplotype is *4.004x2/*5. In the supplemental documentation for Cyrius [4] (Supplement Table S5), the call for this dataset is marked as inconsistent. On that same table, the two parent samples are listed with diplotypes *1/*4x2 (NA19920) and *4/*5 (NA19921), which would not produce a child with *4/*4 without a *de novo* event. Additionally, the column labeled "correct phasing" has "*4x2/*5", indicating that this is a known error in the Cyrius diplotypes. Based on this evidence, we believe the *4.004x2/*5 call produced by StarPhase is correct. A visual representation of the StarPhase graph for this dataset can be seen in Figure 5.

##### 2.4.3 *DPYD*

In Section 2.3, we describe how PharmCAT will allow *DPYD* to generate haplotypes that are not explicitly defined in the database by combining them. Under most circumstances, this simply generates haplotype combinations that StarPhase will not, leading to "NO_MATCH" calls in the StarPhase outputs. However, there are also situations where PharmCAT will *incorrectly* interpret the phase information inside the VCF, creating a potential error in the output diplotype. For example, in HG01891, PharmCAT reports "Reference/[c.85T>C (*9A) + c.3067C>A]", indicating that the two variants are in *cis*. In contrast, StarPhase reports "c.3067C>A/c.85T>C (*9A)", indicating that the same variants were detected but reported in *trans*. Both tools are detecting the same pair of variants, but with alternate phasing. We have copied the relevant parts of the VCF file below (partially truncated for space):

[6]HG02492 public dataset: `s3://1000genomes/1000G_2504_high_coverage/additional_698_related/data/ERR3989019/HG02492.final.cram`

```
chr1 97078987 . G T 61.6 PASS . GT:GQ:DP:AD:PS 0|1:60:45:28,17:96997594
chr1 97883329 . A G 66.2 PASS . GT:GQ:DP:AD:PS 0|1:64:43:19,24:97710720
```

Upon manual inspection, we identified the source of the discrepancy. Inside the VCF file, the two variants have identical genotype entries of "0|1", which would normally indicate two variants that are in *cis* with each other. However, they do *not* share the same phase set ID (PS tag), indicating that the two variants are not in the same phase block and cannot be accurately labeled as in *cis* or *trans* from the VCF. StarPhase correctly recognizes that the phasing of these two variants relative to each other is unknown, and reports the only viable pair of *defined* alleles in the database. If we allow for variant combinations that are not defined (see Section 2.3), then reporting both the *cis* and *trans* options would be correct (i.e., the phasing is unknown). In short, PharmCAT is incorrectly interpreting the phase set information and reporting one of two viable options, while StarPhase is limiting the options to defined haplotypes and reporting the other of the two viable options. The issue with PharmCAT phase interpretation has been reported to the developers[7].

##### 2.4.4 *HLA-B*

For NA19320, StarPhase reported *57:02:01:01/*57:02:01:03 compared to *57:02/*57:03 from our DRAGEN-based comparator. We inspected the consensus sequences generated by StarPhase and did not identify any obvious errors in the sequences (i.e., the reads seemingly support the two sequences). Additionally, the database *57:02:01:01 was an exact match to our generated consensus, while the database *57:02:01:03 had one 1 intronic difference to our other generated consensus. We selected the 4-field *57:03:01:01 haplotype to be representative of the alternate call, and inspected the haplotype using the StarPhase IGV visualizations for *HLA-B*. We identified two variant positions that seemingly distinguish *57:02 from *57:03 and that are within exons (chr6:31354129A>G and chr6:31356247C>A), neither of which were identified in **any** of the HiFi reads. We suspect this is an error in the comparator set, and speculate that it may be caused by the relative homozygosity of the sample.

##### 2.4.5 *MT-RNR1*

For HG02132, StarPhase reports a "Reference" haplotype whereas the DRAGEN-based comparator reported "896A>G". When we manually inspected the alignment, we identified 5 out of 166 reads with the alternate "896A>G" variant. The low frequency in this specific dataset led to a homozygous reference call in the DeepVariant VCF file that is provided to StarPhase, and ultimately the "Reference" haplotype from StarPhase. Since this was reported in the comparator set and detected at low frequency in our dataset, we suspect that this variant is in a heteroplasmic state in this particular sample. We do not consider this an error in either StarPhase or the comparator. However, users should be aware of the difficulties and variability when assessing and reporting heteroplasmic haplotypes from the mitochondria. Future versions of StarPhase may alter our processing for mitochondrial haplotypes, allowing for reporting of low-level heteroplasmy within a dataset.

#### 3 GeT-RM comparison details

GeT-RM samples are commonly used to benchmark PGx end-to-end solutions. In our WGS accuracy collection, we only had three datasets with matching GeT-RM PGx calls (HG00423, NA19317, and NA20509). However, we did have access to an additional 22 datasets that were sequenced on the Twist Alliance Long-Read PGx Panel [8], bring us to a total of 25 HiFi datasets with comparison GeT-RM calls. These datasets were processed using the recommended StarPhase settings for targeted sequencing (adds `--infer-connections --normalize-d6-only` to the command) which help compensate for shorter read lengths and coverage biases when computing the *CYP2D6* diplotype. This section describes the analysis we performed on this data collection.

[7]The PharmCAT phase interpretation issue is being tracked here: https://github.com/PharmGKB/PharmCAT/issues/175

[8]Twist Alliance Long-Read PGx Panel: https://www.twistbioscience.com/products/ngs/Long-Read-Sequencing-Panels

#### 3.1 GeT-RM comparison set

We downloaded the "Consolidated Table of all GeT-RM Pharmacogenetic Reference Material Genotypes" from the CDC website[9]. For our data collections, the vast majority of the comparison calls were derived from a single GeT-RM publication [18], which is labeled as "Reference 2" in the table. This publication includes a significant amount of additional information regarding how the calls were generated, limitations of the calling, sources of ambiguity, and also the list of possible haplotypes that could be included. We highlight specific portions of these limitations in Section 3.2 where we discuss mismatches between the StarPhase calls and this GeT-RM comparison set.

#### 3.2 Discrepancies with GeT-RM

The following sections describe each non-"match" label and how we programmatically identified them. We also include summaries of the identified mismatches below. Detailed comparison information can be found in the Supplementary Tables (described in Section 5.2).

##### 3.2.1 Parsing errors

Parsing errors were caused by unusual diplotype strings in the GeT-RM collection. Typically, these seemed to capture ambiguity of some form, although the exact nature of each ambiguous diplotype was sometimes unclear. There were only two detected parsing errors, both in *CYP2B6*. Manual inspection deemed them as a likely match, although the ambiguity makes this result unclear.

##### 3.2.2 Not tested in comparator

The reference for the majority of the GeT-RM calls was [18]. Table 1 from [18] includes a list of all tested loci and alleles for each of the genes in the collection. We identified many alleles that were called by StarPhase but that were not a part of this test set. These calls were labeled as "Not tested in comparator", and we added notes to our Supplementary Table indicating which of the haplotypes were not included in the comparator test set. This category is the closest match to the "New DB haplotype" category from the HPRC comparison, but it is broader because some of these haplotypes may have been known and simply not validated for the tests that were used to build the comparator set.

##### 3.2.3 Absent from StarPhase DB

There was one haplotype, `UGT1A1`*60, which was completely absent from the StarPhase database but present in the comparator set. We verified that our upstream data source (CPIC) was also missing this haplotype in the haplotype definitions for *UGT1A1* and also that PharmCAT is also not reporting this haplotype. This haplotype was retired in March 2017 after a publication showing that *60 has normal function [15]. We note that other authors have identified this issue when evaluating tools on GeT-RM calls for *UGT1A1* as well [12].

##### 3.2.4 Mismatches

We identified 7 total mismatches between StarPhase calls and the GeT-RM comparator, which are categorized below:

1. Five "NO_MATCH" diplotypes - StarPhase reported a "NO_MATCH" diplotype for five diplotypes, indicating that the collection of observed variants do not form two *exact* matching haplotypes from the CPIC database. Four of these are from *DPYD*, which frequently has this issue as described in Sections 2.3 and 2.4.3. For *DPYD*, this happens whenever two or more variants are found on a single haplotype. The last "NO_MATCH" call was from NA19226 *CYP2B6*. One haplotype seemingly supports *6 and the other seemingly supports a combination haplotype, [*18 + *20]. For all five, our Supplementary

[9]GeT-RM PGx table: `https://www.cdc.gov/labquality/get-rm/inherited-genetic-diseases-pharmacogenetics/pharmacogenetics.html`

Table reports each identified variant in the notes field. In each case, the variants from GeT-RM were identified, but additional variants identified by StarPhase led to the final "NO_MATCH" call.

2. One likely phasing error in the comparator set - For NA20509 *CYP2B6*, GeT-RM reports *1/*7 whereas StarPhase reported *5/*6. We note that the difference in these two diplotypes is whether two variants are found in *cis* (*1/*7) or *trans* (*5/*6). Manual inspection of the aligned BAM file clearly supported that the two variants were found in *trans*. Additionally, the authors of the GeT-RM comparator set noted this exact possibility: "When c.516G>T, c.785A>G, and c.1459C>T are present together, the most likely interpretation is *1/*7 with all of the variants in *cis*, although *5/*6 in *trans* cannot be ruled out" [18]. Given this evidence, we suspect that the true diplotype for NA20509 *CYP2B6* should be *5/*6 as reported by StarPhase.

3. One unexplained discrepancy - One final discrepancy was identified with HG01190 *CYP2C9*. GeT-RM has the diplotype as *2/*61 whereas StarPhase called *1/*61. We manually inspected the aligned BAM file and noted that the rs1799853 was found in a heterozygous state in our dataset (27 reads supporting REF, 28 read supporting ALT). The ALT allele for rs1799853 is found in both *2 and *61, implying that our data does not support a *2/*61 haplotype since one haplotype must have the REF allele for rs1799853. While we cannot definitively rule out a sequencing or data processing issue upstream of StarPhase, the coverage of this region is relatively high and strongly supportive of a heterozygous call. We suspect this is instead some error in the GeT-RM comparator set, though we do not have an indisputable explanation for the source of this error.

### 4 Additional analyses

#### 4.1 Analysis of trio datasets

We performed a trio-based analysis using a collection of 37 internal whole-genome sequencing trio datasets. For each trio, we ran StarPhase on the child and both parents. We then categorized each trio as "consistent" if the child haplotypes could be explained through normal inheritance patterns from the parent haplotypes. For most genes, this means one haplotype from the maternal line and one from the paternal. We included special inheritance cases for *G6PD* on chrX (single-copy for males) and *MT-RNR1* (resides on the mitochondria and should be inherited through maternal line). The summary results for this trio comparison are in Figure 2.

Of the 777 comparison made through the trio evaluation, we identified 752 that were consistent with expected inheritance patterns. We note that 16 genes were fully consistent, meaning that all comparisons for that gene were consistent. There were 22 comparisons where the child and/or a parent had a "NO_MATCH" call. As noted earlier, these missing calls occur when a haplotype is detected that is not present in the CPIC database (see Section 2.3). As with previous comparisons, *DPYD* is the gene where this problem manifests the most.

Lastly, we found 3 total inconsistent inheritance patterns. Of these, one occurs in *DPYD* and is seemingly caused by two variants that are inherited in *cis* but reported in *trans* due to database limitations. The other two are both in *MT-RNR1* and seemingly caused by low coverage in the maternal dataset, leading to drop-out of variants in the VCF file that was provided to StarPhase. We include a de-identified table with details on each specific comparison including the exact diplotype calls reported by StarPhase for each dataset in each trio in our Supplemental Tables (described in Section 5.3).

#### 4.2 HLA consensus delta metrics

*HLA-A* and *HLA-B* are the two most diverse genes that StarPhase diplotypes, and haplotypes for these genes are defined by cDNA and DNA sequences. StarPhase generates full-length consensus sequences for each reported haplotype, providing the edit distance in basepairs from the consensus to the reported haplotypes. We collected these edit distance metrics for both cDNA and DNA, and Figure 3 shows the distributions within our cohort.

The vast majority of cDNA sequences for both genes *exactly* match their corresponding database sequence (overall, ≈99% for both *HLA-A* and *HLA-B*). The majority of DNA sequences also *exactly* matched their

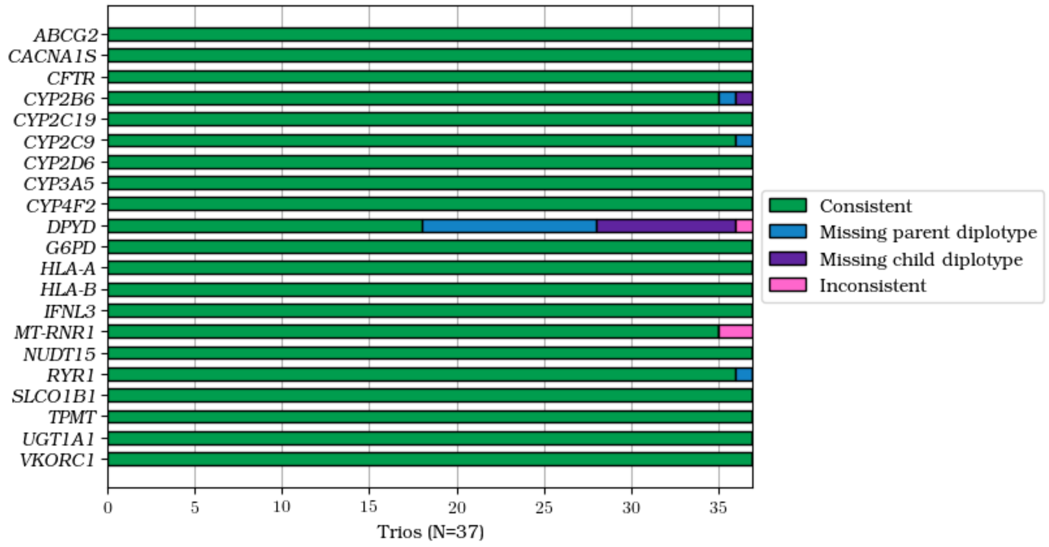

Figure 2: Summarized trio consistency. This figure shows a summary of the trio consistency analysis performed on 37 internal whole-genome sequence trios across all 21 genes reported by StarPhase. Of the 777 trio evaluations, we identified 752 that were consistent with expected inheritance patterns (accounting for exceptions in *G6PD* and *MT-RNR1*). 16 of the 21 genes were full consistent, meaning that all comparisons for the gene were consistent. There were 22 where the child and/or parent diplotype call could not be described with existing database definitions, the majority of which are from *DPYD*. Lastly, we identified three inconsistent inheritance patterns. The one from *DPYD* is seemingly caused by two variants in *cis*, which is not captured by the CPIC database. The last two are seemingly caused by low coverage in the maternal dataset along a portion of the mitochondria, leading to dropout of variants in the maternal VCF.

database sequences, but not as many as the cDNA (respectively, ≈95% and ≈92%). This matches our expectations for two reasons. First, we expect changes to the cDNA to be rarer due to conservation of function (i.e., changes are more likely deleterious), so capturing the relative diversity is likely easier in the shorter cDNA. Second, historically, the focus for HLA has been on 2-field annotations due to functional impact, so many other assays do not capture DNA sequences leading to under-representation (or absence) of DNA sequences in IMGT/HLA.

For sequences that did not exactly match the database, most of the DNA haplotypes differed by exactly one basepair (respectively, ≈4% and ≈6% of total). Through manual inspection of a subset of these "Off-by-one" haplotypes, we identified some that are likely true differences relative to the database, whereas others are more likely consensus and/or sequencing errors. The majority of these likely errors were indels located in homopolymer regions, which are known to be enriched for indel errors from sequencing. We also identified a smaller percentage of haplotypes that had two or more differences between the consensus sequence and the database DNA sequence (respectively, ≈1% and ≈2% of total). After manually inspecting a subset, we believe many of these are candidate undescribed haplotypes relative to the IMGT/HLA database (i.e., a novel 4-field haplotype; see Supplemental Materials). Lastly, a small fraction (most observable as ≈0.5% of EAS population) of the haplotypes did not have a DNA sequence we could compare against in the IMGT/HLA database (i.e., cDNA only). Both the absent DNA sequences and those identified with differences relative to the existing database are candidates for improving the IMGT/HLA database.

#### 4.3 Cohort zygosity distribution

The population diplotypes were each labeled as heterozygous or homozygous and then collapsed by gene (ancestry was ignored). *G6PD*, which is located on chrX, has the extra hemizygous category for males. *MT-RNR1*, which is located on the mitochondria, uses homoplasmic and heteroplasmic labels. We include a category for the "NO_MATCH" calls from StarPhase, which occur when the observed diplotype does not

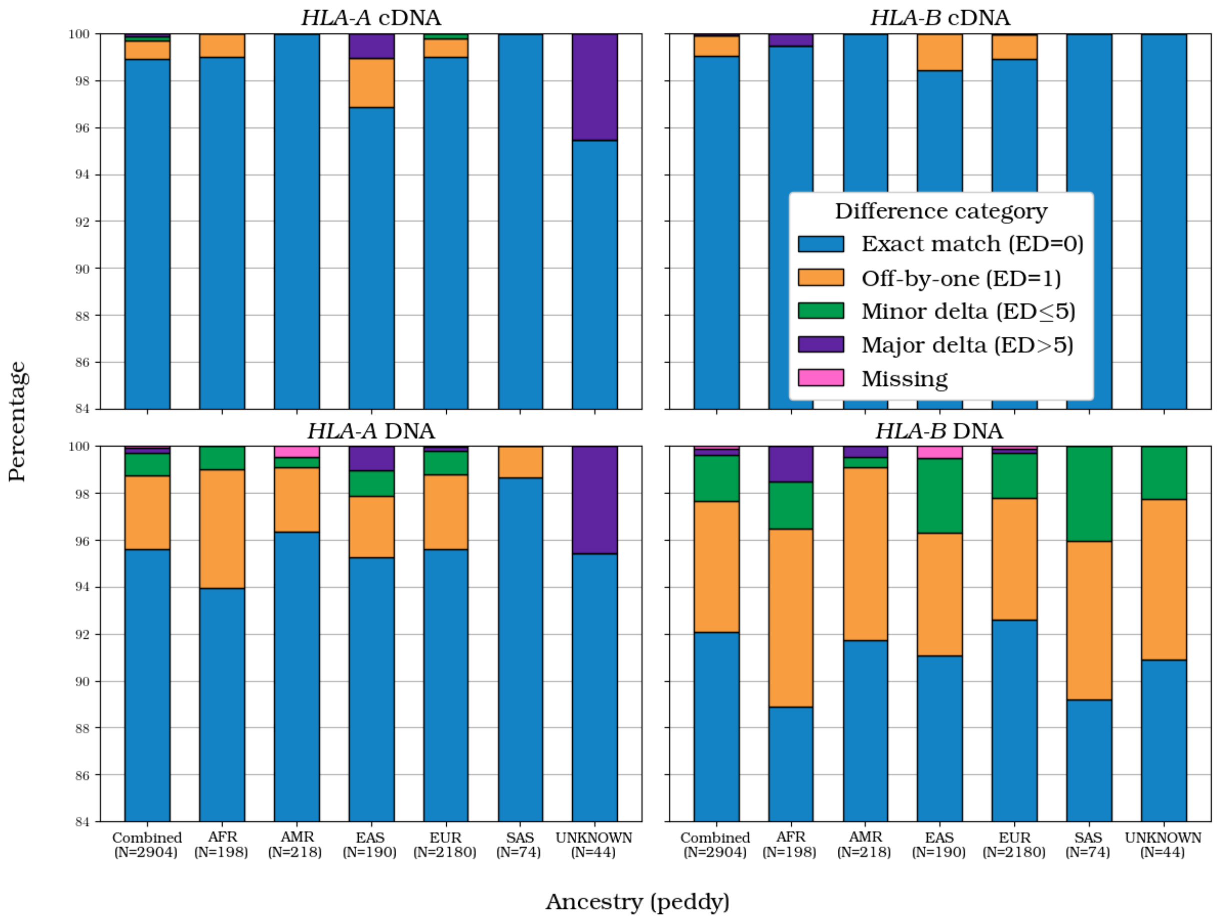

Figure 3: HLA gene summaries. These figures show summary edit distance metrics for *HLA-A* (left) and *HLA-B* (right) by computed ancestry in our cohort. The top row shows the edit distance between the consensus cDNA sequences from StarPhase and the corresponding cDNA sequence from the IMGT/HLA database. We note that the vast majority of these exactly match the sequence from the database. The bottom row shows the edit distance between the consensus DNA sequences from StarPhase and the corresponding DNA sequences from the IMGT/HLA database. We see an elevated number of differences in the DNA sequences relative to the cDNA sequences, but the vast majority still exactly match. "Off-by-one" indicates a single basepair difference between the consensus and the database, some of which are likely caused by sequencing and/or consensus errors. Minor deltas and especially major deltas are more likely undescribed haplotypes relative to the IMGT/HLA database. "Missing" indicates that no DNA sequence is available in the IMGT/HLA database for comparison, making them good candidates for updating the database.

match database definitions. Lastly, ambiguity in the final diplotype was filtered at each data collection site, leading to a reduced total count for some genes. This was rare with one exception: *UGT1A1*, where ≈15% of diplotypes were ambiguous in our cohort. The ambiguous diplotypes for *UGT1A1* all came from one contributing site which had older sequencing datasets with shorter read lengths, leading to more frequent incomplete phase blocks for the gene. We did not observe this issue in any other contributing site. Figure 4 shows the cohort zygosity distribution for each gene.

### 5 Supplemental Tables explanations

This section contains details on the accompanying Supplemental Tables that are bundled in a single Excel spreadsheet file (filename `starphase_sup_tables.xlsx`). These tables are packaged separately from this

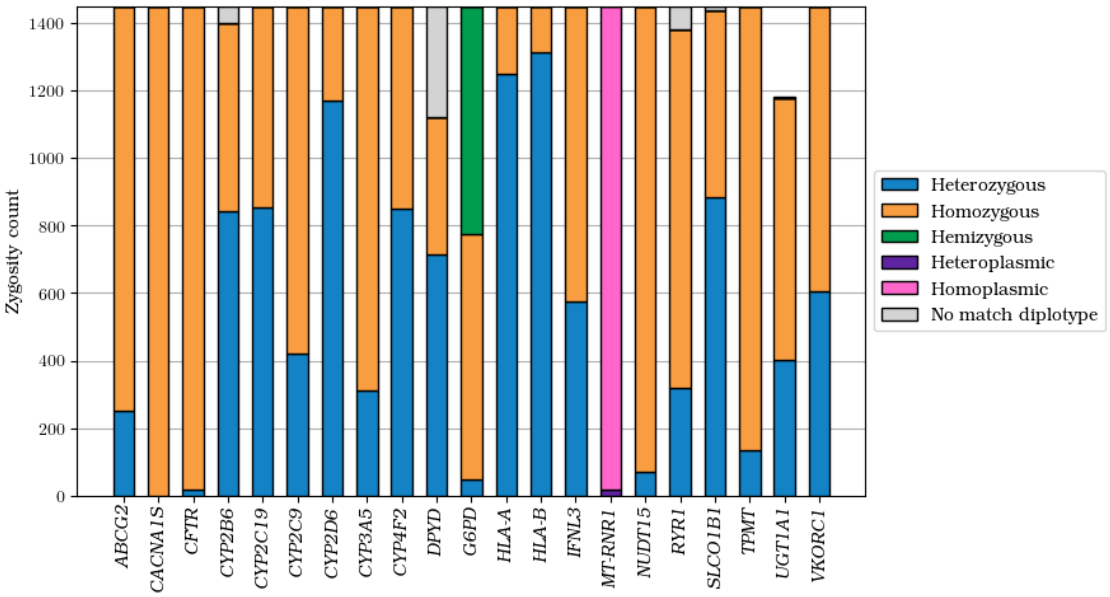

Figure 4: Zygosity summary per gene in our population cohort. The overall zygosity counts for each gene was totaled, ignoring ancestry. Most genes had only heterozygous and homozygous categories. *G6PD*, which is located on chrX, had an additional category for males. *MT-RNR1*, which is located on the mitochondria, was labeled with heteroplasmic or homoplasmic instead. An additional label for "NO_MATCH" diplotypes is also counted. Lastly, some diplotypes were ambiguous, meaning two diplotypes were equally likely based on the observations. These are excluded at the collection sites, leading to reduced total zygosity counts for some genes. This most obviously impacts *UGT1A1*.

document to make them more easily accessible to researchers and developers looking to leverage the data programmatically.

#### 5.1 HPRC comparison table – `1_comparator_set_detailed`

This table contains the full collection of StarPhase diplotype calls as well as the comparator call. We include a column for classification of the StarPhase call relative to the comparator as well as detailed notes on each different diplotype. The following is a description of each comparison label:

1. `mismatch` - Indicates that the two diplotypes are different, and cannot be automatically explained by known discrepancy sources. Each mismatch was manually inspected and notes are provided in the table and in Section 2.
2. `reporting_discrepancy` - Indicates that the sample has a haplotype that is not described by the CPIC database. StarPhase reports these as a "NO_MATCH" diplotype whereas the comparator will sometimes generate a haplotype that does not exist in the database. See Section 2.3 for more details.
3. `new_db_haplotype` - Indicates that StarPhase produced a haplotype that was not part of the Cyrius call set, limited to *CYP2D6*.
4. Matches - Multiple labels are broadly categorized as a match in the main document.
   (a) `match` - Indicates that the result exactly matched. For HLA genes, this means a full 4-field match. For *CYP2D6*, this comparison is limited to core alleles due to comparator limitations.
   (b) `third_field_match` - For HLA genes, this means a 4-field comparator was available but the StarPhase diplotype only matched through the 3rd field. Each was manually inspected and notes are provided in the table and in Section 2.

(c) `second_field_refinement` - For HLA genes, this means a 2-field comparator was available and StarPhase 2-field matched that and refined it with a full 4-field haplotype.

5. `no_comparator` - Indicates that a suitable comparator was not identified for this dataset. We provide these as a reference point, but caution users as we have not performed any orthogonal validation of these additional diplotype calls.

#### 5.2 GeT-RM comparison table – `2_getrm_detailed`

This table contains the full collection of StarPhase diplotype calls as well as the comparator GeT-RM diplotype. We include a column for classification of the StarPhase call relative to the comparator as well as detailed notes on each different diplotype. The following is a description of each comparison label:

1. `different` - Indicates that the two diplotypes are different, and cannot be automatically explained by known discrepancy sources. Each difference was manually inspected and notes are provided in the table and in Section 2.

2. `parsing_error` - Indicates that the comparator was not trivially parsable by machine, and that the two diplotypes were manually inspected. Notes are provided for each of these, and both were deemed a "Match".

3. `absent_starphase_db` - *UGT1A1**60 was retired from the CPIC database in March 2017. Each entry with this label includes at least one *60 haplotype from GeT-RM, and could not be cleanly compared. Given the high agreement with HPRC comparators for *UGT1A1*, we suspect that these are correct, updated diplotypes for the GeT-RM datasets.

4. `not_tested_comparator` - Indicates that StarPhase produced a haplotype that was not part of the GeT-RM tested haplotypes (see [18]). In some cases, these are haplotypes that were not defined when the comparator was generated, and in other cases the haplotypes were defined but simply not captured by the various assays. Give the high agreement with HPRC comparators for these genes, we suspect that these are correct, updated diplotypes for the GeT-RM datasets.

5. `matches` - Indicates that the result exactly matched. For *CYP2D6*, this comparison is limited to core alleles due to comparator limitations. We note that some automatic filtering and translating of the GeT-RM diplotypes was automatically performed to generate a match. For example, the *VKORC1* "G" from GeT-RM translates into "rs9923231 reference (C)" by modern CPIC haplotype definitions.

6. `no_comparator` - Indicates that a comparator was not available through GeT-RM for this dataset. We provide these as a reference point, but caution users as we have not performed any orthogonal validation of these additional diplotype calls.

#### 5.3 Trio comparison table – `3_trios_detailed`

This table contains the full collection of StarPhase trio diplotypes used for the trio analysis. These samples have been stripped of identifying information, so no sample labels are present. We include a column for classification of the trio and notes for discrepancies.

#### 5.4 Population diplotype table – `4_bulk_diplotypes`

This table contains the bulk diplotype metrics for our entire cohort. Samples were grouped by ancestry and sex, and then the observed gene-diplotype counts for each grouping is reported as a row.

#### 5.5 Population haplotype/metric table – `5_bulk_haplotypes`

This table contains the bulk diplotype metrics for our entire cohort. Samples were grouped by ancestry, and then the observed gene-haplotypes counts for each ancestry is reported as a row. Autosomal genes received two haplotype counts per diplotype, genes on chrX (*G6PD*) received either 1 count for male datasets or 2 counts for female datasets, and genes on the mitochondria (*RNR1*) received 1 count per dataset with discrepant calls flagged as "heteroplasmy". Additional metrics that were reported in this document or the main document are also included for ease-of-use (e.g., HLA delta metrics for cDNA and DNA and *CYP2D6* functional categories).

### 6 Methods details

#### 6.1 Summary

This section provides a summary of how the underlying methods work within StarPhase. These methods are grouped by the genes they are applied to. The CPIC genes are all diplotyped using just a VCF file, whereas the HLA genes and *CYP2D6* require an aligned read file.

##### 6.1.1 CPIC genes

Broadly speaking, the variant calls in each of the 18 CPIC genes are relatively "clean". The variant calls inside these genes typically do not have complications caused by sequence homology, low-complexity sequence, or other genomic features that may cause issues during the alignment and variant calling process. As a result, StarPhase will implicitly trust the variant information provided by the input VCF file for each of these genes, relying on the upstream tooling to correctly identify and phase variants.

However, the representation of variants in the VCF file tends to differ from the representation in the CPIC database [19]. For example, a VCF file is typically normalized (mean extra bases are trimmed) and left-aligned with an anchor base. In contrast, there are many entries in the CPIC database that do not follow this convention. There are entries where extra bases are provided, entries that do not provide full length representations of reference or alternate sequences, and entries that are neither left-aligned nor anchored to a particular fixed base in the reference genome. Thus in StarPhase, the bulk of the compute for these genes is just normalizing both the VCF and CPIC variants such that matching variants can be cleanly identified. Additionally, StarPhase will store any available phase information, including whether a variant is phased at all and which phase set ID that variant is a part of. Importantly, StarPhase will correctly interpret variants that are phased but in different phase sets, meaning they are effectively unphased relative to each other. In contrast, PharmCAT seemingly ignores phase set IDs, which can lead to errors in the haplotype assignments. This is the source of all four mismatch errors between StarPhase and PharmCAT in the *DPYD* gene (see Section 2.4.3). Lastly, if no VCF entry is identified that matches the database variant, then the variant is assumed to be homozygous for the reference allele.

Once all variants are identified with their phasing information, StarPhase creates all possible combinations of those variants into haplotype pairs. If phasing is provided, then variants that are in cis or trans will have that order enforced during pair generation. As noted above, StarPhase respects phase set IDs here, meaning that even phased VCF files may have multiple combinations to check if the gene spans multiple phase blocks. For each haplotype pair, StarPhase compares the result to the list of defined haplotypes from the CPIC database. If *both* haplotypes in the pair *exactly* match a defined haplotype, then the pair is one candidate diplotype. Importantly, multiple candidate diplotypes are possible when ambiguity in the phasing is present (this is most likely when unphased VCF files are used). In the event of multiple candidate diplotypes, all are reported in the output (this is the same behavior as PharmCAT). In the event that no pair has two exact haplotype matches, StarPhase will report `NO_MATCH/NO_MATCH` for the diplotype.

##### 6.1.2 HLA genes

In contrast to the CPIC genes, *HLA-A* and *HLA-B* do not have trustworthy variant calls when using most standard workflows. This is primarily caused by other homologous HLA genes that incorrectly map in the reference HLA locus, creating false positive and false negative variant calls throughout the region.

Additionally, the IMGT/HLA database [1] is a sequence-oriented database, meaning that instead of storing variant differences between a haplotype and some reference, it instead stores full-length cDNA and/or DNA sequences for each haplotype. Of note, these haplotype sequences are not equally represented in the database. During development we identified 1) haplotypes without a DNA sequence, 2) haplotypes without a cDNA sequence (very few, these are ignored in StarPhase), 3) haplotypes that only contain part of the cDNA sequence (e.g., exons 2-4), and 4) haplotypes where the DNA sequence cover different lengths of reference genome. This combination of imprecision in variant calling as well as imbalance in database representation required an alternate approach to diplotyping these HLA genes.

The approach taken for *HLA-A* and *HLA-B* is to first generate two consensus sequences and then label those sequences based on the IMGT/HLA database. To generate the two consensus sequences, StarPhase identifies all reads that fully span the gene region. These reads are then re-aligned to the reference gene region, tracking the edit distance from the mapping to the reference region. If this value is too high, the read is discarded as it most likely is a mis-mapping of another HLA gene. For each read that is kept, the sub-string that mapped to the reference gene region is extracted. Additionally, a sequence representing the spliced cDNA is also extracted for each kept read. Finally, these sub-strings are run through the dual consensus algorithm (see Section 6.2.2) to generate up to two full-length haplotypes.

Conceptually, the dual consensus algorithm is searching for an optimal split of the read observations into two groups, each one representing a parental haplotype. First, it searches for two distinct cDNA sequences by running dual consensus on the spliced cDNA sequences. If that only finds one consensus group (i.e., the two haplotypes are identical at the cDNA level), StarPhase re-runs dual consensus but with the full-length DNA sequences, potentially identifying differences at the DNA level. Once the consensus groupings are identified, StarPhase runs the single consensus algorithm on each group's DNA sequences separately, generating a representative full-length haplotype DNA sequence for each group. Additionally, these full-length DNA sequences are spliced into a representative cDNA sequence for database matching.

Finally, each cDNA/DNA haplotype sequence is scored against the database. In the database, HLA sequences are labeled such that the third field represents matching cDNA and the fourth field represents matching DNA, thus matching to cDNA is prioritized over matching to DNA. One nuance of the IMGT/HLA database is that not all haplotypes are equally represented within the database. In particular, the cDNA and DNA sequences may be missing or only contain a portion of the region (e.g., only a subset of exons or different amount of surrounding DNA sequence). To handle this nuance, StarPhase aligns each database haplotype to the consensus sequence to generate a total edit distance for the consensus, tracking where each edit is relative to the consensus. Then, when comparing two candidate haplotypes to determine which matches the consensus better, it will first evaluate them only within regions that are overlapping in the consensus alignments, effectively excluding an edit that may only occur in one sequence because it happens to be longer in the database (see Figure 1 for an example where this is relevant). In the event of a tie (which is rare), the full sequences will be used to score it on the edit distance ratio (i.e., i.e., $edit\_distance/haplotype\_length$). For both comparisons, we first score on the cDNA alignments and then secondarily on the DNA alignments. In contrast to the CPIC gene approach, this approach will report the best matching haplotype even if it is not an exact match to the database sequence. Note, there are debug options available in StarPhase to report the full-length consensus sequences in a FASTA format, which may be of interest to researchers investigating potentially novel haplotype sequences such as the one in Figure 1. The labeled haplotype pair is reported as the final diplotype for each HLA gene.

##### 6.1.3 *CYP2D6*

Similar to the HLA genes, *CYP2D6* often does not have trustworthy variant calls, but for different reasons. First, *CYP2D6* has relatively common deletion (*5) and duplication (x2, x3, or xN) haplotypes. Second, there is a homologous pseudogene, *CYP2D7*, nearby that can cause mis-mappings of reads within the locus. Finally, there are known hybrid alleles between *CYP2D6* and *CYP2D7* that start similar to one gene and then switch to look more like the other gene, with *68 and *36 being the most common hybrid alleles. This creates a problem where there are two haplotypes in an individual that may have anywhere from zero to three copies of *CYP2D6* on each haplotype. Biologically, more than three copies may be possible, but there are no annotations for those haplotypes in PharmVar. Additionally, these copies may be completely identical or slightly different within or between haplotypes within an individual dataset. In StarPhase, we break this problem into two core components: 1) identifying each allele that is present in the dataset and generating a consensus sequence for it, and 2) "chaining" these alleles together into an optimal diplotype (a pair of haplotypes) that best explains the observations (i.e., reads) in the dataset.

StarPhase solves the first part of this problem by using the multi-consensus approach described in greater detail in Section 6.2.3. At a high level, we identify regions of interest within each read, extract their sequences, and then build a collection of consensuses from them. However, there are some nuances to how we utilize this algorithm for *CYP2D6*. First, there are multiple "types" of regions that we identify through this approach, and each type is effectively treated as its own multi-consensus problem. Each type has a defined reference region in the StarPhase configuration file. These types are grouped based on homology for multi-consensus and are enumerated here: 1) *CYP2D6*, *CYP2D7*, and hybrids; 2) REP6 region; 3) REP7 region; 4) *CYP2D6* deletion signature (*5); 5) the "spacer" region; and 6) the "link_region", which is the section of DNA commonly found between *CYP2D6* and REP7. We refer the reader to the PharmVar Structural Variation in *CYP2D6* document[10] for a visual representation of how these regions are expected to be connected.

The second nuance is that the consensus algorithm can over-split groups in regions with higher error rates. In particular, we identified homopolymer errors as a common problem when using this multi-consensus approach. To resolve this issue, the multi-consensus algorithm first operates on a homopolymer-compressed representation of each sequence. Then, a secondary multi-consensus splits the groups further based on the full-length sequences (see Section 6.2.4 for details). Next, each group is assigned a label from the database corresponding to either the type (e.g., REP6) or the full-length allele (e.g., CYP2D6*4.001). Finally, StarPhase will merge two groups back together if A) they share the same homopolymer compressed consensus and B) they have identical labels. This allows StarPhase to identify false splits caused by homopolymer errors without losing true splits caused by variants that get masked by homopolymer compression and correspond to different alleles in the database (e.g., *2 and *41 are indistinguishable when homopolymer compressed).

The second step of this algorithm is to "chain" these consensuses together into two haplotypes, much like an assembly. To do this, StarPhase takes each read and scores all of the identified regions of interest against each consensus sequence. These scores are based on edit distance from the read to the consensus, where lower scores are better. Each read has an "optimal" chain of regions associated with it (i.e., a best scoring chain), although there can be ambiguity when a read starts or stops in the middle of a region of interest. With whole-genome HiFi sequencing, we commonly observe many reads with four or more of these regions linked together, allowing for a strong baseline signal to link all of the identified alleles. These optimal chains are used to create an internal graph representation, that is then traversed to create candidate chains. Figure 5 shows an example graph created by StarPhase for debugging purposes with a loop due to the presence of a *4x2 haplotype. This loop creates an ambiguity what the optimal haplotype is, leading to StarPhase generating three possible candidate haplotypes (*4, *4x2, and *4x3) that will be scored in conjunction with other candidate haplotypes from the graph.

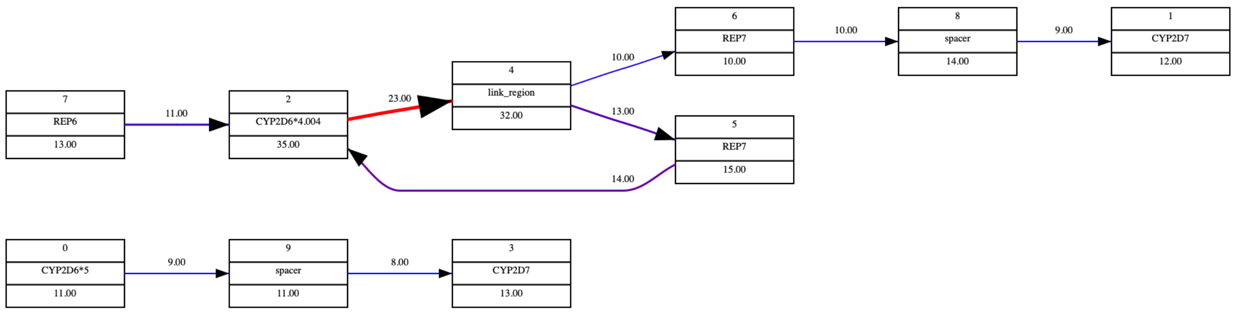

Figure 5: Example *CYP2D6* graph. This figure shows an example *CYP2D6* graph generated on a dataset with *4x2/*5. The top section contains a *4 allele with evidence looping back to *4 before continuing to a *CYP2D7* allele, and the bottom section shows a distinct haplotype from *5 through a second *CYP2D7* allele.

[10]PharmVar Structural Variation document: `https://a.storyblok.com/f/70677/x/d842ef4108/cyp2d6_structural-variation_v3-1.pdf`

In practice, most graphs are composed of two distinct paths that can be trivially solved by eye or by StarPhase. However, some haplotypes or diplotype combinations are more challenging to resolve: 1) haplotypes with exact duplications can create ambiguous loops (see Figure 5); 2) haplotypes with hybrids can create ambiguous links (e.g., *68 (hybrid allele) can be easily confused with *4 (hybrid source allele) and *CYP2D7*); and 3) identical alleles on separate haplotypes can create ambiguity in copy number within the dataset (e.g., a dataset that is *4/*4x2). To address these complexities/ambiguities, StarPhase scores all possible haplotype pairs using a combination of metrics based on the read observations:

1. Edit distance penalty - For a given haplotype pair, each read is placed optimally along the pair to identify the place with the least edit distance between the read and the consensuses. Then, this placement is compared to the "optimal" placement (i.e., the minimum edit distance for this read to any consensus), and the delta is applied as a positive penalty. This penalty is basically penalizing any haplotype combinations that do not fully explain the direct observed allelic sequences. This penalty is, by far, the dominant penalty in the scoring system and most optimal diplotypes will have a near-0.0 score here, indicating that they are one of the best possible explanations for the observed data.

2. Multi-nomial likelihood coverage penalty - For a given haplotype pair, StarPhase can count how many times each allele occurs within the pair and assign a likelihood of the read count distribution given that pair. For example, for a *1/*4x2 diplotype, we would expect roughly double the number of reads for *4 relative to *1. Thus, observing 15 reads for both *1 and *4 would have a lower likelihood than observing 10 reads for *1 and 20 reads for *4. This penalty is mostly for tie-breaking haplotype combinations that can fully explain the reads but are ambiguous due to possible duplications in the haplotypes.

3. Copy-number penalty - However, we do not want the algorithm to allow for uncontrolled copy number, so there is an additional penalty whenever an extra copy of an allele is incorporated. This penalty acts as cost that must be overcome by the above observational or multi-nomial penalties in order to report a duplication event.

4. Unexpected penalty - There are certain duplication and hybrid combinations that are known and expected to be found together in the human population. For example, *68 is almost always paired with *4, *36 is almost always paired with *10, and only certain alleles have been found in duplicated states. Whenever a haplotype pair has unexpected allele combinations, this penalty is increased to represent that the combination is not something that has been previously observed in the human population.

5. Inference penalty (only when inference is enabled) - For whole-genome sequencing, inferring connections between alleles in the graph is not necessary due to direct observational evidence from long reads. However, targeted sequencing datasets will often lack any direct observations of connecting alleles due to shorter reads. In those instances, StarPhase has an option that allows the algorithm to infer a connection if it has been previously established in the population (e.g., the expected pairs described above). For example, if we observe *1, *4, and *68 alleles, it is much more likely for *4 and *68 to be on one haplotype than any other combination. However, each inference comes with a penalty to prevent over-inference of the allele connections.

Each of the above penalties have been tuned to operate together to identify the most likely combination of haplotypes. Once all haplotype pairs have been scored using the above system, the one with the lowest score (i.e., fewest penalties) is reported as the optimal *CYP2D6* diplotype for the dataset.

#### 6.2 Consensus generation

This section describes the core consensus generation algorithms that are the most computationally expensive part of StarPhase. At a high level, these algorithms are all based on using a dynamic wavefront algorithm (DWFA) to both track optimal alignments to a collection of strings and nominate extension characters to create an optimal consensus. The first section describes how DWFA works to generate a single consensus sequence, and the following sections describe how we extended that into a dual consensus (one or two consensus sequences) and then multi-consensus ($N >= 1$ consensus sequences). These algorithms are bundled as part of the `waffle_con` crate we have made available on GitHub[11].

##### 6.2.1 Dynamic WFA consensus

Underlying all of the consensus algorithms is an adaptation of the wavefront algorithm (WFA) [13] that allows for dynamic selection of extension characters. In brief, WFA is a pairwise alignment algorithm that achieves increased efficiency by taking advantage of regions of high similarity between two sequences. It does this by operating along a "wavefront", which is a collection of maximally extended diagonals if you allows for a particular maximum "cost". In our application, we define this cost as the edit distance, and note that the wavefront with edit distance $(i + 1)$ can be calculated from the wavefront with edit distance $i$. If the actual edit distance between two sequences of length $n$ is $s$, then the algorithm runs in $O(ns)$ time and uses $O(s^2)$ memory [13]. Thus, for highly similar sequences (i.e., $s$ is small), this algorithm is very efficient even for long sequences. This is a key feature of the algorithm that is advantageous when using long, high accuracy HiFi reads.

We adapt this method into a dynamic WFA (DWFA) algorithm. With DWFA, one sequence (an observed read sequence) is static, serving as an anchor for the algorithm. The other sequence is instead generated dynamically at run-time. Conceptually, this requires little changes to the WFA algorithm other than the ability to pause while waiting for the next character to get generated dynamically. However, while the algorithm is paused, the state of the DWFA algorithm can be observed, allowing us to view where the wavefronts are relative to the static sequence. This observational process allows us to *nominate* characters that would extend the "other" sequence *without* increasing the edit distance parameter. If we think of a simple case where we do not know the static sequence, *but* we do have the DWFA for it, a naive algorithm could simply 1) request a nominating character, 2) extend the "other" sequence with that character, and 3) repeat (1) and (2) until no more nominees are left. This would basically copy the static sequence into our "other" sequence.

The above example operates on a single sequence, effectively copying it into our "other" sequence. The DWFA consensus extends this by operating on multiple static sequences (i.e., the observed read collection) while generating a shared "other" sequence that is ultimately the consensus for the collection. Thus, the core loop is 1) each DWFA nominates one or more characters, 2) the character with the highest number of votes is used to extend our consensus, and 3) repeat (1) and (2) until no more nominees are left. At the end of this process, we have a consensus sequence that theoretically has the smallest combined edit distance to the entire observed read collection.

Of course, there are situations where there are multiple possible extension characters, usually due to true variation (e.g., a SNP) or errors in sequencing. Typically, these can be trivially resolved by simply picking the character with the most votes as the next character in the consensus. However, there are situations where this is sub-optimal (especially if true variation is involved), so our implementation uses a search tree that allows the DWFA algorithm to split into multiple possible paths if the criteria is met. As the algorithm progresses, we can accumulate many possible extension paths, most of which are inactive due to relatively high edit distance with low progression into the read collection. Our implementation includes heuristics to prune these inactive paths, typically leaving a handful of active candidate consensus sequences. We refer the reader to the GitHub repository for greater understanding of the nuances and details of how the DWFA consensus algorithm works in practice.

##### 6.2.2 Dual consensus

Given the above DWFA consensus algorithm, we extend the approach to generate either one or two consensus sequences from a single sequence collection. In practice, our primary use case for this is clustering and building consensus sequences for the HLA genes. Recall that a search tree implementation runs in conjunction with the DWFA processes to navigate the consensus search space. The dual consensus algorithm simply extends this search tree to allow for either one or two active consensuses at a time. If only one consensus is active (which is how the algorithm is initialized), then the algorithm operates very similar to the main DWFA consensus approach. However, when multiple possible extension characters are identified, we use

[11] `waffle_con` crate: `https://github.com/PacificBiosciences/waffle_con`

various heuristics to determine whether the characters are a possible divergence point between two consensus sequences. If so, the "node" that gets generated in the search tree will be flagged as a dual node with two active consensus sequences. Additionally, the edit distance cost for that node will be the sum of the minimum edit distance between each read sequence and *either* consensus sequence. Of course, for homozygous diplotypes, the algorithm heuristics are designed to prevent dual nodes from getting generated from sequencing artifacts.

When a dual node is initially generated, the distance between the two candidate consensuses is exactly 1. As more extensions are made, this difference will grow and "sort" the input sequence collection into two groups that match each consensus. Once a sequence from our collection is clearly matching one consensus over the other, we stop tracking the worse match and assume that the sequence is part of the better matching consensus. For the HLA loci, the high sequence divergence typically leads to clean splits of the read sequences into two groups, each of which will ultimately form one consensus haplotype. We refer the reader to the GitHub repository for greater understanding of the nuances and details of how the DWFA dual-consensus algorithm works in practice.

##### 6.2.3 Multi-consensus

While the dual consensus approach works quite well for the HLA loci, we needed the ability to generate multiple consensus sequences for the *CYP2D6* gene. For example, a dataset with diplotype "*68+*4/*36+*10" is a very reasonable possibility from the population. This dataset would have four distinct alleles present in the diplotype, and the possibility of two highly similar *CYP2D7* genes nearby. This could create a situation with up to six potential consensus sequences that are all mixed into the read collection.

Conceptually, the dual consensus algorithm is splitting the sequence collection into two groups of sequences such that the similarity within a group is higher than the similarity between the groups. In other words, it is clustering the sequences into two groups. Multi-consensus simply extends this clustering by calling dual-consensus repeatedly until the cluster returns a single consensus (i.e., it does not find a valid split based on the heuristics). This approach can return anywhere from 1 to $N$ consensus clusters, where $N$ is the number of input sequences in the collection. In practice, there are many heuristic constraints in place to prevent over- or under-clustering of the sequences into consensus groups. We refer the reader to the GitHub repository for greater understanding of the nuances and details of how the DWFA multi-consensus algorithm works in practice.

##### 6.2.4 Priority consensus

Lastly, we encountered problems with the *CYP2D6* locus where the multi-consensus algorithm would over-split the clusters into multiple due to sequencing errors in homopolymer regions. This would create 2+ consensuses that were completely identical except for the addition/subtraction of a base in one or more homopolymer regions. To address the issue, we created an adaptation of the multi-consensus algorithm that operates on multiple versions of the same input sequence. In particular, we provided the homopolymer-compressed (HPC) version of the sequence and also the full-length version. The algorithm would first cluster the sequences based on the HPC version, and then provide a secondary clustering based on the full-length sequence. Thus, the output would be a group of primary clusters as well as one or more secondary sub-clusters of each primary cluster. Section 6.1.3 describes how the *CYP2D6* diplotyping step used these sub-clusters to identify the final clusterings. We refer the reader to the GitHub repository for greater understanding of the nuances and details of how the DWFA priority-consensus algorithm works in practice.
